# Switching environments, synchronous sex, and the evolution of mating types

**DOI:** 10.1101/2020.07.31.230482

**Authors:** Ernesto Berríos-Caro, Tobias Galla, George W. A. Constable

## Abstract

While facultative sex is common in sexually reproducing species, for reasons of tractability most mathematical models assume that such sex is asynchronous in the population. In this paper, we develop a model of switching environments to instead capture the effect of an entire population transitioning synchronously between sexual and asexual modes of reproduction. We use this model to investigate the evolution of the number of self-incompatible mating types in finite populations, which empirically can range from two to thousands. When environmental switching is fast, we recover the results of earlier studies that implicitly assumed populations were engaged in asynchronous sexual reproduction. However when the environment switches slowly, we see deviations from previous asynchronous theory, including a lower number of mating types at equilibrium and bimodality in the stationary distribution of mating types. We provide analytic approximations for both the fast and slow switching regimes, as well as a numerical scheme based on the Kolmogorov equations for the system to quickly evaluate the model dynamics at intermediate parameters. Our approach exploits properties of integer partitions in number theory. We also demonstrate how additional biological processes such as selective sweeps can be accounted for in this switching environment framework, showing that beneficial mutations can further erode mating type diversity in synchronous facultatively sexual populations.

## I. INTRODUCTION

Evolution is a fundamentally noisy affair [1]. It is therefore no surprise that over the last century theorists have increasingly sought to mathematically understand the effects of randomness on evolutionary models. Such noise has many distinct forms. The foundations of mathematical population genetics are rooted in models that capture how genetic drift (demographic noise), emerging from uncertainty in the order of birth and death events in finite populations, can drive population dynamics [2]. Meanwhile a more ecologically-oriented approach has been to consider the noise that might arise from uncertainty in environmental conditions [3, 4] (environmental noise), with a particular emphasis in the evolutionary literature on transitions between discrete environmental states [5, 6] (capturing, for instance, an organism’s switching behavioural responses to the fluctuating environment). However, in the last decade in particular, there has been an increasing interest in mathematically understanding the dynamics of populations subject to both demographic noise and environmental switching [7–10].

Beyond simply presenting a mathematical challenge, developing analytic techniques to attack such systems is important for understanding a host of problems in biology. One simple yet acute example is that of a population switching between environments in which selection is present in one environment and absent in the other [11]. Here one must understand the interplay of quasi-deterministic dynamics on the one hand (in the selective regime) and entirely noisy dynamics in the other (where genetic drift dominates). In this paper we will consider just such an evolutionary problem, demonstrating how it can be modelled and, more importantly analysed, quantitatively.

Mating types are self-incompatible gamete classes that can be understood as ancestral forms of the more familiar sperm-egg system [12, 13]. Unlike populations with true sexes however (which are defined by the size dimorphism between their gametes) the number of mating types (which are morphologically similar) is not restricted to two [14]. Instead, mating types are expected to experience negative frequency dependent selection, with rare types favoured due to their increased opportunities for finding a compatible mate of a distinct, non-self mating type. This has an important dynamical consequence; novel mutant mating types, which are initially rare, should nearly always successfully establish within a resident population, and therefore the number of mating types in a species is predicted to increase through time [15]. However, while species with many mating types are possible rising to many thousands in some fungi [16], species with more than 10 are rare and most have just two [17, 18]. This disagreement between simple evolutionary reasoning and empirical evidence sets the stage for a classic evolutionary paradox [19].

Although many theories have been proposed to explain this discrepancy (reviewed in [19]), most rely on a deterministic selective advantage for two mating types, such as increased mating success between two types in pheromone signaling and receiving roles [20], or decreased cytoplasmic conflict between two types in donor-receiver organelle inheritance roles [21]. In contrast, [15] demonstrated that under differing assumptions about the gamete encounter rate dynamics, the strength of selection for more than two types could be reduced. It was then verbally suggested that demographic stochasticity may play a role in further limiting the number of mating types. However without the analytic tools to quantify this effect, the hypothesis that genetic drift could govern mating type number through a balance between mutations and stochastic extinctions was somewhat neglected within the mating type literature, despite being well-established in the related but distinct system of self-incompatibility alleles in plants [22].

More recently, simulations were used to show that in a population that switched between sexual and asexual environments, mating type extinctions became more likely, with negative frequency dependent selection absent in the asexual regime, and the population dynamics entirely dominated by genetic drift [23]. Extending this logic, [24, 25] showed analytically that an increased rate of asexual to sexual reproduction would lower the number of mating types expected under a mutation extinction balance, and indeed that available empirical data showed a positive correlation between the rate of sexual reproduction and the number of mating types in these species.

For mathematical simplicity these latter models [24, 25] considered sexual reproduction to be occurring asynchronously, i.e., with each reproductive event having a fixed probability of sexual vs asexual reproduction. However this simplification fails to capture a biologically relevant aspect of reproduction in these species; sexual reproduction tends to be triggered by changing environmental conditions [26], such as falling nutrient levels [27] or other stress cues [28], and is thus synchronized in time across the entire population [29]. With such dynamics better captured by a switching environment model, it is interesting to ask what quantitative differences this increased level of biological realism might generate. More importantly, this shift in modelling framework also enables us to explore a richer array of biological questions.

In addition to demographic stochasticity, selective sweeps have been suggested as a mechanism that may increase mating type extinction rate and, therefore, further limit the number of mating types [30]. However the effect of these selective sweeps on mating type number can only be seen in asexual environments, where beneficial mutations are linked to the mating type background on which they arise [31]. In a sexual or even partially sexual environment, genetic recombination breaks down associations between beneficial mutations and mating types, allowing the mutations to spread through the population without distorting mating type frequencies. Quantifying the effect of selective sweeps on these dynamics therefore requires a shift in modelling approach, away from simplified mathematical assumptions of asynchronisity and towards more biologically realistic switching environments [30]. In this paper we focus on this problem, describing a modelling framework and developing a mathematical analysis suitable for the task.

This article is organized as follows: in Section II, we present the switching-environments model in which a population transition between entirely sexual and entirely asexual reproductive modes. Section III is dedicated to studying how the distribution of mating types changes as a function of the switching and mutation rates. We focus in particular on the regimes of fast, slow and intermediate environmental switching. In Section IV we demonstrate how this new modelling framework allows us to address the issue of selective sweeps. Finally, we present the conclusions in Section V.

## II. MODEL DEFINITIONS

### A. Population dynamics

We consider a population genetics model similar to that proposed in [24]. The model describes a population of *N* individuals, who each are of a particular mating type. The different mating types are labelled by the index *i*. The population follows a dynamics similar to the Moran model (i.e., coupled birth-death events in continuous time) but now allowing three possible types of events: asexual reproduction, sexual reproduction, and mutation. Each reproduction event implies the removal of one individual so that the size of the population remains constant. Mutation events imply the introduction of a new mating type. We write *n*_*i*_ for the number of individuals of mating type *i*, and *M* for the total number of mating types in the population. These are time-dependent quantities. The number of mating types ranges from *M* = 1 (all individuals are of the same type) to *M* = *N* (each individual is of a different mating type).

In the model by Constable and Kokko in [24] both sexual and asexual reproduction were possible at any time, each occurring with a fixed rate. In this paper we instead consider the more biologically realistic scenario of a population that engages in sexual reproduction in response to changing environmental conditions. While the environment itself may be described by a continuous quantity (such as temperature) the population’s response to the environment is binary (whether to engage in sexual reproduction or not). We therefore develop a model that switches stochastically between two different states, denoted *S* and *A*, respectively. We write *σ ∈ {S, A}* for the environmental state. When the environment is in state *S*, only sexual reproduction is possible, and when it is in state *A*, only asexual reproduction is possible. We assume that the switching between these two environmental states occurs independently of the composition of the population, with rate *λ*_*S→A*_ from *S* to *A*, and *λ*_*A→S*_ for switches from state *A* to *S*. The switching processes can be written as

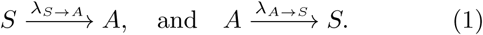

In the sexual environmental *S*, we assume that any pair of individuals can reproduce, provided they belong to two different mating types. For example, one parent may be of mating type *i*, and the other parent of any other non-*i* mating type. The probability that this occurs for two individuals sampled at random from the population is *n*_*i*_(*N− n_i_*)*/N* ^2^. The offspring inherits the mating type of either parent with equal probability 1*/*2. To keep the population size fixed, another individual (type *j*) is simultaneously chosen uniformly at random to die. The rate for events in which an offspring of type *i* is generated and an individual of type *j* removed from the population is then

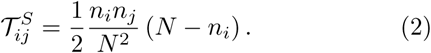

We express time in units of generations, so that there are order *N* events in the population per unit time. This means that rate in Eq. (2) has an extra factor *N* in comparison to the rate used in [24].

In the asexual environmental *A*, reproduction follows the standard neutral Moran model. One individual is chosen uniformly at random to reproduce, and the offspring inherits the mating type of the parent. As above, another individual is simultaneously chosen at random to die. The rate for events in which an individual of type *i* reproduces and an individual of type *j* is removed is then given by

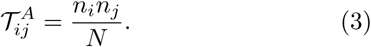

Following [24], we describe mutations as events in which one individual changes to a new mating type not currently present in the population. This leads to the rate

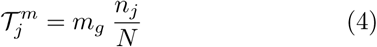

for mutation events from type *j* to a new type. The parameter *m*_*g*_ sets the typical number of mutations per generation in the population. The raw mutation rate is given by *m* = *m*_*g*_/*N*. Defined in this way, Eq. (4) can also be interpreted biologically as capturing migration events from a highly diverse mating type pool.

The dynamics of the model above are summarised in Figure 1. We will refer to this as the ‘full model’ in the remainder of the paper. It describes a Markovian process. At each point in time its state is described by the state of the environment (*S* or *A*), and by the state vector of the population, **n** = (*n*_1_, *n*_2_, *…*). The *i*-th entry in this vector indicates how many individuals of mating type *i* are present in the population. We have ∑_*i*_ *n*_*i*_ = *N*, and the number of non-zero entries in **n** indicates the number of mating types currently present in the population.

**FIG. 1.**
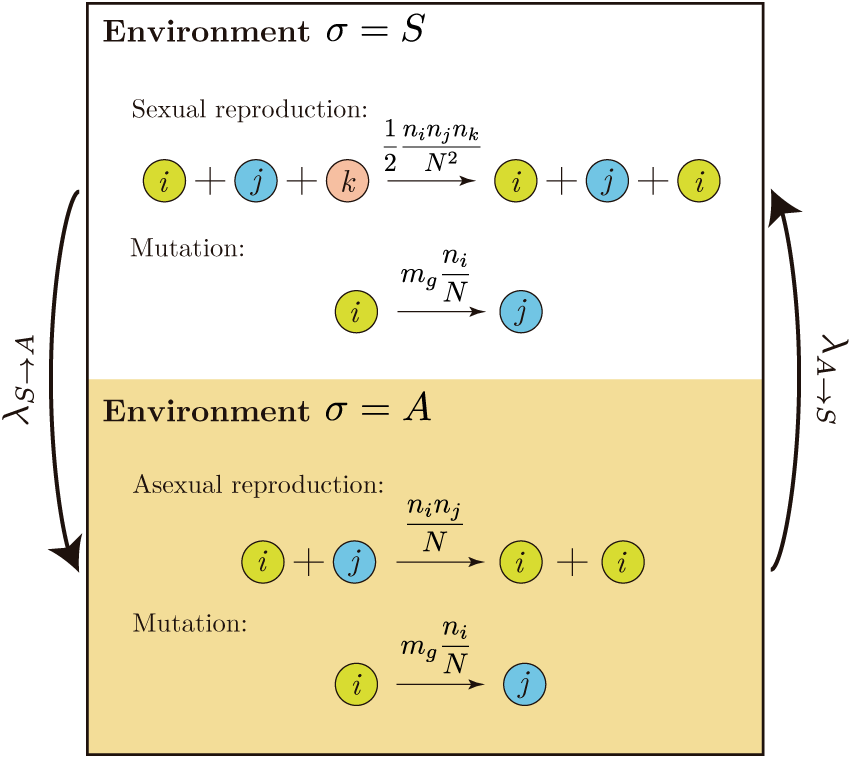
Illustratition of the full model and the events occurring in the population: sexual reproduction, asexual reproduction, and mutation. In each of these events one individual is replaced by another of a different mating type. Transitions between environments occur independently of the state of the system at rate *λ*_*S→A*_ from *σ* = *S* to *σ* = *A*, and *λ*_*A→S*_ from *σ* = *A* to *σ* = *S*.

### B. Environmental dynamics

Given that the environmental switching is independent of the composition of the population, the long-time probabilities to find either environmental state can be written down straight away,

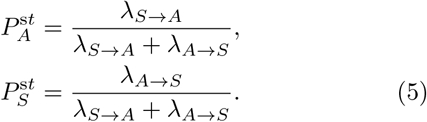

To ease the notation, we write 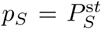 for the probability to find the environment in state *S*. This indicates the rate of sexual reproduction.

The average time the environment spends in each of the two states between switches is given by

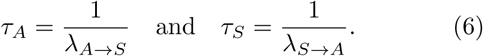

The average time to switch from one state to the other and back, is then

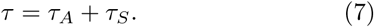

### C. Reduced model

In order to analyse the dynamics of the population, we will focus on a *reduced model*, describing only the number of mating types *M*. This number changes over time through the birth-death and mutation events in the population. In a birth-death event the number of mating types can decrease by one (if the individual that dies is the last individual belonging to a particular mating type). When a mutation event occurs, the number of mating types in the population increases by one (unless the mutating individual is the last of its type). We are interested in the stochastic process for *M*, and will describe it with effective rates

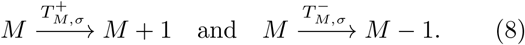

For example, 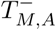 is the rate with which a mating type is driven to extinction when the environment is in state *A*, and when there are currently *M* mating types present in the population. Figure 2 illustrates this approach. The reduced model focuses on the dynamics of the number of mating types *M*, without regard for the numbers *n*_*i*_ of individuals belonging to each mating type. The stochastic process for *M* is of course dependent on the composition of the vector state **n** in the full model, and as such, the reduced model constitutes an approximation of the dynamics in the full model.

**FIG. 2.**
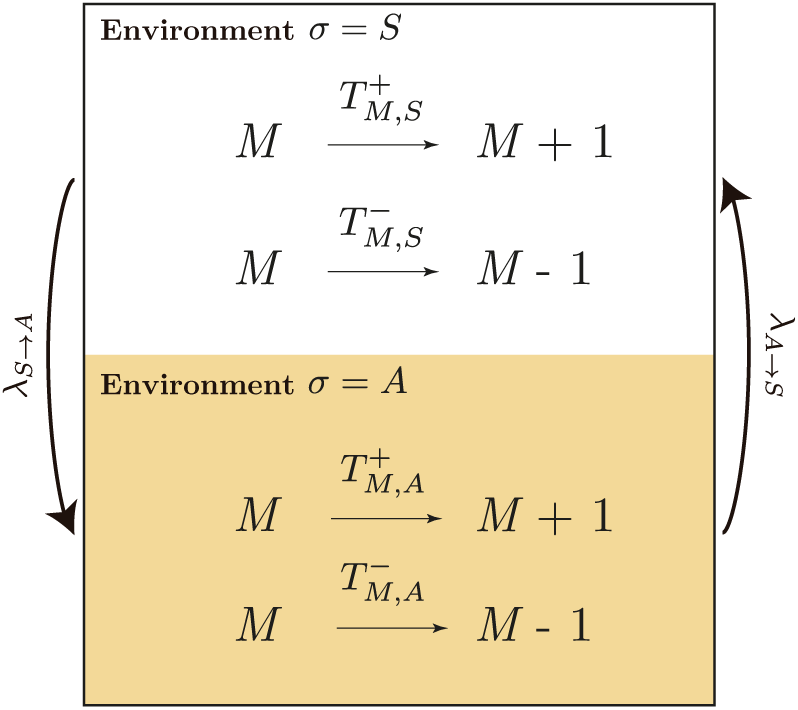
Schematic representation of the *reduced model*, where the focus is on the dynamics of the number of mating types *M* instead of the individuals. This model is described by effective birth and death rates 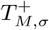 and 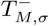, with *σ ∈ {S, A}*.

The analytical challenge is to derive suitable expressions for the 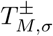. While this is difficult for the case of switching environments, progress can be made by focusing on the case of a fixed environmental state, *σ* = *A* or *σ* = *S*. In this case, the population will tend to a stationary state, described by the joint distribution of the number of mating types, *M*, and the vector of abundances **n**. This distribution can be obtained analytically using an approach similar to that of [24].

We then proceed to use this distribution to calculate the rates 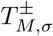 for the dynamics of *M*. To do this, we focus on marginals for specific values of *M* and use methods from number theory [32, 33] to sum over partitions **n** of the *N* individuals into mating types. Importantly, this approach accounts precisely for all possible transitions in which a change on state **n** leads to an increase or decrease in the number of mating types *M*. Our calculation of these rates relies on fixed environmental states *A* or *S*. To make this clear in the notation we write 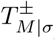 for the rates computed in this way. Further details of the calculation can be found in Sections S1 and S2 of the Supplementary Material. The outcome of this approach is an analytical solution for the rates 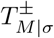. For environment *σ* = *A*, we find

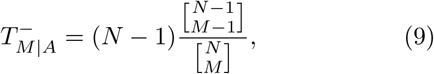

and

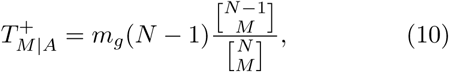

where 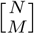 is the unsigned Stirling number of the first kind [34]. For environment *σ* = *S*, the rates become

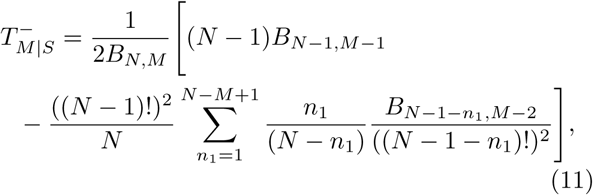

and

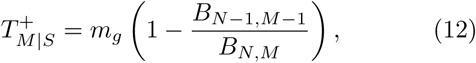

where *B*_*k,l*_ = *B*_*k,l*_(*y*_1_, …, *y*_*k−l*+1_) is the incomplete Bell polynomial [35]. The arguments *y*_*i*_ are the sequence *y*_*i*_ = (*i −* 1)!(*N −* 1)_*i−*1_, with (*N −* 1)_*i−*1_ the falling factorial of (*N −* 1) with respect to (*i −* 1), given by 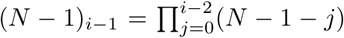. Further details can be found in Section S2 B 4 in the Supplementary Material. In Figure 3 we demonstrate the accuracy of the predictions for the rates 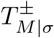 when compared against direct measurements of the rates from simulations of the full model with fixed environmental state.

**FIG. 3.**
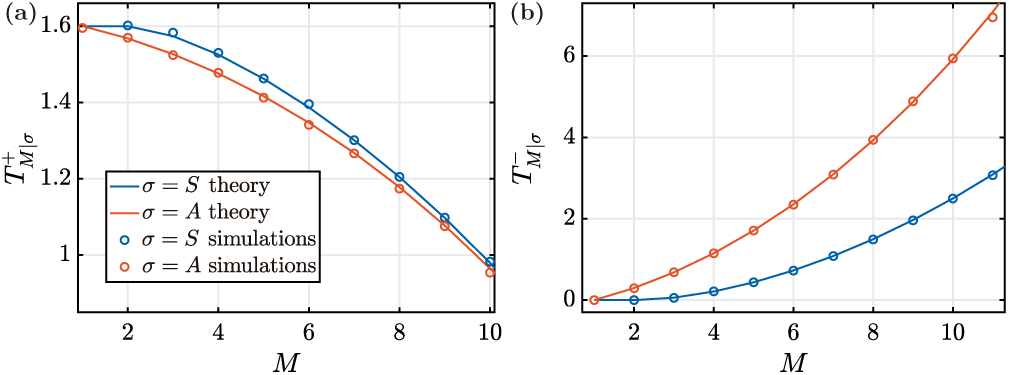
Theoretical predictions of rates 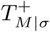 and 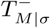 against numerical simulations of the full model with fixed environments, *σ* = *S* and *σ* = *A* respectively. Parameters are *N* = 16 and *m*_*g*_ = 1.6.

We now proceed to discuss the properties of these rates as a function of *M*. We first focus on the rates 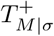, i.e., events in which the number of mating types increases. The introduction of new mating types occurs when one individual mutates from one mating type to another. In order for *M* to increase in this process, the mutating individual must not be the only representative of its type. As a consequence 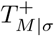 tends to decrease with increasing *M* : if a large number of mating types is present in the population, then it is likely that some of these will only be represented by a small number of individuals, and possibly by a single member of the population. A mutation event involving this individual then does not lead to an increase of the number of mating types.

In birth-death events the number of mating types can decrease, irrespective of whether reproduction is sexual or asexual. A reduction of *M* occurs when the individual that dies in such an event is the only representative of its mating type. Given that the size of the population is fixed, this is more likely to be the case when the number of mating types is large, hence the rate 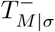 increases with *M*.

Using these rates we can obtain the stationary distribution for the number of mating types under fixed environmental conditions using standard results for continuous-time Markov chains [38]. We have

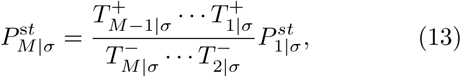

with

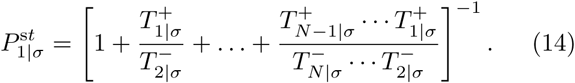

Using this expression, we can write the stationary distribution in closed form for environment *σ* = *A*,

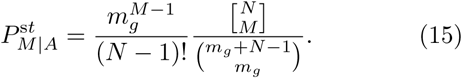

For environment *σ* = *S*, we use Eq. (13) with rates given by Eqs. (11) and (12). In Section S2 D of the Supplementary Material we show that these predictions for 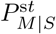 and 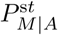 are in good agreement with numerical simulations.

## III. STATIONARY DISTRIBUTION FOR THE NUMBER OF MATING TYPES UNDER ENVIRONMENTAL SWITCHING

While in previous studies [24, 25, 30] the frequency of facultative sexual reproduction was measured by a single parameter (the probability of a reproduction event being sexual), the switching environment model requires two parameters, *λ*_*A→S*_ and *λ*_*S→A*_. Importantly, while the probability of finding the population in a sexual state remains constant for a fixed ratio *λ*_*A→S*_/*λ*_*S→A*_ (see Eq. (5)), the population dynamics qualitatively changes as the values of these parameters are changed. This is illustrated in Figure 4, where typical time courses of the number of mating types present in the population are shown (left panels) along with the corresponding stationary distributions 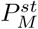 (right panels). We next give a brief overview of the behaviour of the model.

**FIG. 4.**
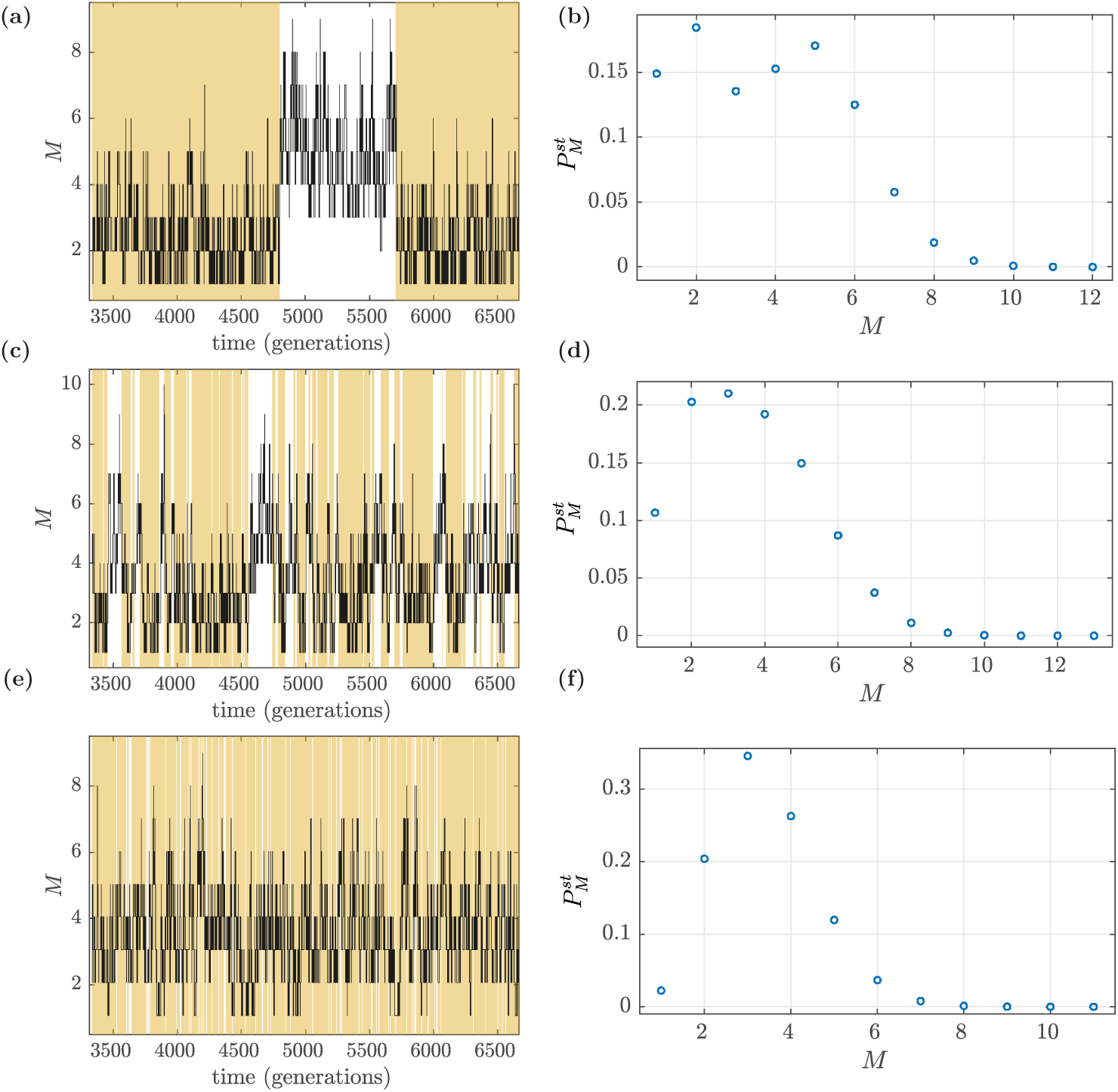
Sample path of the temporal evolution, and stationary distribution of the number of mating types *M*. The environment switches between *σ* = *S* (entirely sexual reproduction) and *σ* = *A* (entirely asexual reproduction). Coloured regions in (a), (c), and (e) represent the *σ* = *A* environment. Data is shown for different switching regime. Panels (a) and (b) illustrate the case of slow switching (*λ*_*A→S*_ = *λ*_*S→A*_ = 10^−5^), panels (c) and (d) of intermediate switching (*λ*_*A→S*_ = *λ*_*S→A*_ = 10^−3^), and panels (e) and (f) of fast switching (*λ*_*A→S*_ = *λ*_*S→A*_ = 10^−1^). Simulations have been carried out by using the Gillespie algorithm [36, 37] in the full model. The stationary distributions shown in panels (b), (d), and (f) have been obtained by time-averaging a long run until *t* = 10^7^, with a time *t* = 10^5^ left to equilibrate. Parameters used: *N* = 30 and *m*_*g*_ = 0.3.

In general we see in Figure 4 that while the number of mating types fluctuates, the number is typically higher when reproduction is sexual. In the sexual environment, rare mating types experience a reproductive advantage, with their their per capita reproductive rate proportional to (*N − n_i_*) (see Eq. (2)). This means that novel mutants (or migrants) establish in the population with high probability [39]. In contrast, in the asexual environment mutants have no particular advantage as all mating types reproduce with the same per capita rate (see Eq. (3)). In this case, given sufficient time, mating types are driven to extinction as a result of neutral genetic drift.

Figure 4 shows three different regimes of environmental switching. In the upper panels the environment is slow compared to the typical time scales of the population dynamics. The stationary distribution of the number of mating types can then be bi-modal, as shown in panel (b). Here mating type numbers greater than *M* = 1 in the asexual regime are only maintained by mutation (or migration) providing a supply of new types. The distribution of *M* becomes unimodal when the typical time scale of environmental switching becomes comparable to the time scale of the evolutionary process in the population (panels (c) and (d)), and it remains unimodal when the environment is much faster than the population dynamics (panels (e) and (f)).

In the following sections we seek to quantify these dynamics mathematically. We begin by considering the limits of slow and fast environmental switching (Figure 4 (a,b) and (e,f), respectively), as these prove analytically tractable. We then go on to consider the range of intermediate switching. In order to illustrate the accuracy of our approximations as compared to simulations, we will use parameters compatible with manageable computing time (e.g. low population sizes and high mutation rates, for which the dynamics more rapidly approach a stationary distribution). In Section III C 4 we will explore more biologically reasonable parameter regimes that are prohibitively expensive to investigate through simulation alone.

### A. Slow environmental switching

When the environmental switching is slow (Figure 4 (a) and (b)) the system spends sufficient time in each environment for the number of mating types to reach stationarity. One then expects the overall distribution of the number of mating types 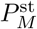 to be the weighted average of the stationary distributions from each environment. Mathematically, this means

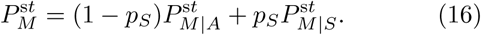

This result can be obtained analytically from the master equation of the system (see below in Eq. (22)) in the limit of slowly switching environments, see Section S3 D 1 in the Supplementary Material. This approximation was also used in [7] for a game theory model with switching payoff matrices. The probability 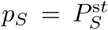 is given in Eq. (5), while the probability 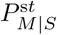 and 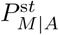 are the stationary distributions for *M* assuming that the environmental state is fixed to *S* or *A*, respectively. These are given in Eqs. (13) and (14).

In Figure 5 (a) and (b), we compare this prediction for the limit of slow environmental change against numerical simulations. We show the distribution 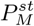 as function of the fraction of time *p*_*S*_ spent in the sexual environment.

**FIG. 5.**
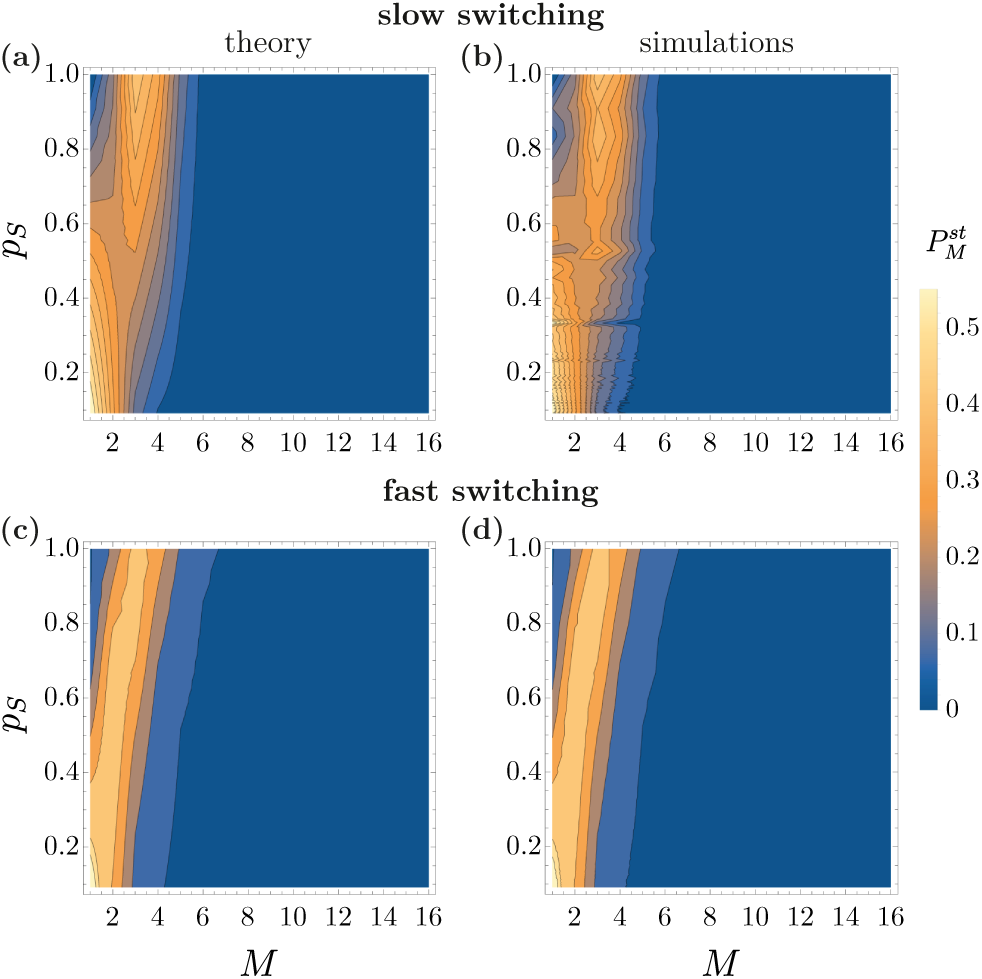
Stationary distribution 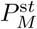 as function of *M* and *p*_*S*_, for slow switching regimes (upper row) and fast switching regimes (lower row). Panels (a) and (c) show the result obtained from numerical simulations for parameters *N* = 16, *m*_*g*_ = 0.16 with *λ*_*A→S*_ = 10^−6^ (for (a)) and *λ*_*A→S*_ = 10^3^ (for (c)). Panels (b) and (d) show the corresponding theoretical predictions in Sections III A and III B, respectively.

The upper panels in Figure 6 illustrate the behaviour of 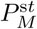 in the regime of slowly varying environments. In particular we show how this stationary distribution changes with the rate *p*_*S*_. We have now chosen a larger population than in Figure 5, and we show results for different mutation (or migration) rates *m*_*g*_. As seen in the figure, the distribution is unimodal if the environment is predominantly in one of its two states (i.e., *p*_*S*_ is close to zero or one). The distribution is bimodal when the environment spends similar fractions of time in each state (*p*_*S*_ *≈* 1*/*2), independent of the rate at which new mating types are added. We also see that the mode of the distribution for 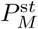 is lower if reproduction is always asexual (*p*_*S*_ = 0) than in the case of obligately sexual reproduction (*p*_*S*_ = 1). As *m*_*g*_ increases (new mating types arrive more frequently), the distribution gets wider around its peak, and the most probable number of mating types is shifted to higher values. Naturally, for very high (and biologically unrealistic) values of *m*_*g*_ the mode of 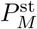 will be equal to the population size *N*.

**FIG. 6.**
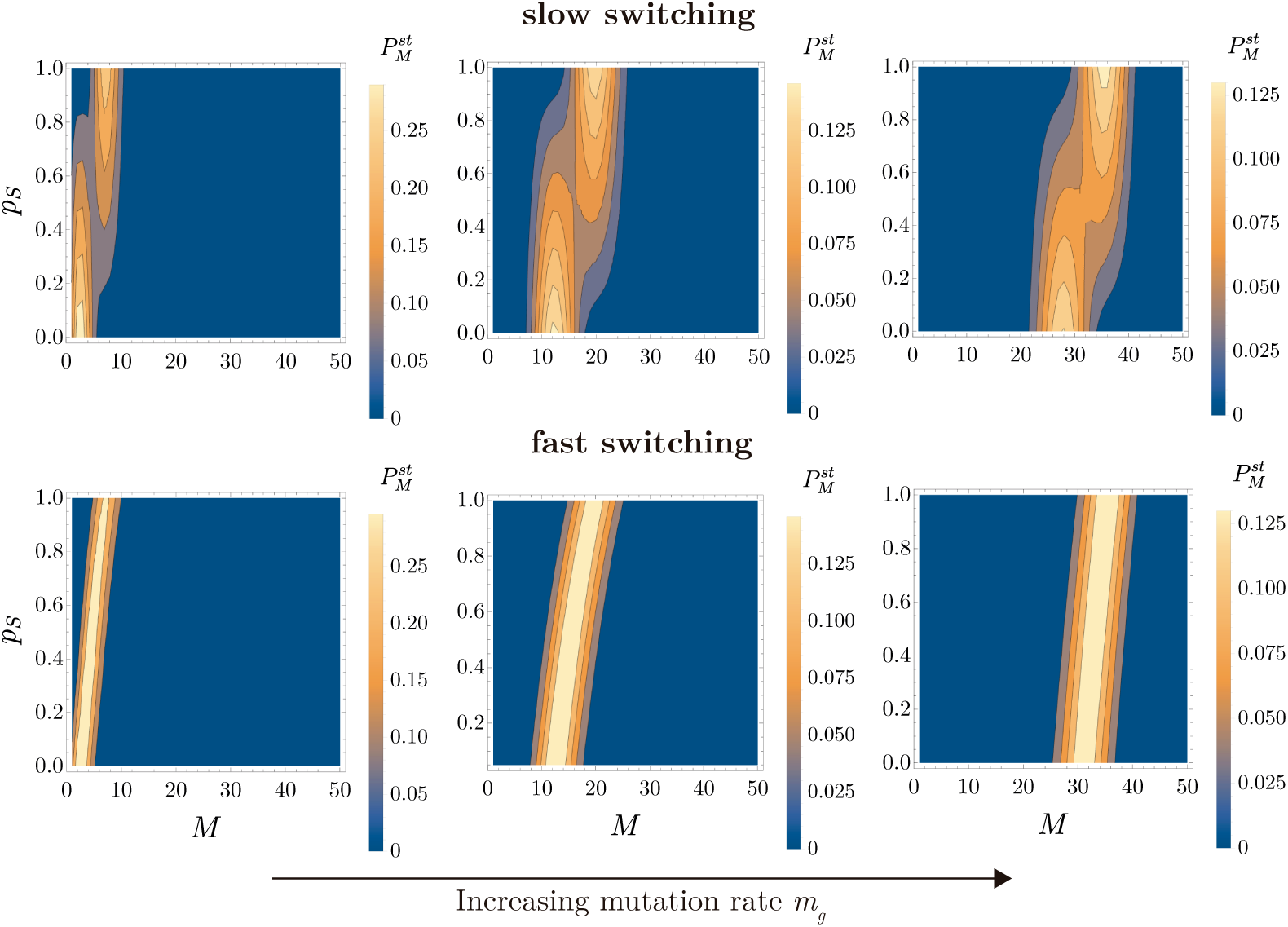
Theoretical prediction for stationary distribution 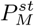, for slow switching (upper row), and fast switching (lower row) as function of the number of mating types (*M*) and the average fraction of time spent in the sexual environment, *p*_*S*_. The theoretical predictions for the two switching regimes are calculated from Eqs. (16) and (18), respectively. Panels (a) and (d) show the case of low mutation rate (*m*_*g*_ = 0.5), panels (b) and (e) are for intermediate mutation rate (*m*_*g*_ = 5), and panels (c) and (f) for high mutation rate (*m*_*g*_ = 50). Population size is *N* = 50.

### B. Fast environmental switching

The simulation data in Figure 4 (e) and (f) illustrates the behaviour of the population in the limit of very fast environmental switching. Unlike in the regime of slow- switching environments, the stationary distribution 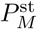 then only exhibits one peak.

To estimate the 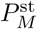 in this regime, we follow the analytical approach developed in [7] for game theoretic models and calculate weighted averages of the transition rates 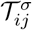 for all pairs *i, j*,

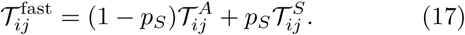

This leads to

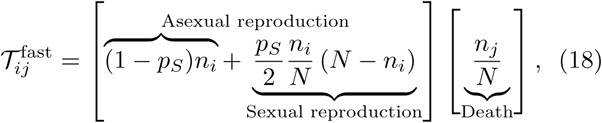

and one recovers the limit studied in [24] for the case of asynchronous facultative sex. This is a model with a constant environment with an effective sex rate *p*_*S*_. This sex rate determines the effective birth and death rates, 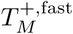 and 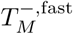 Further details of the derivation of these rates are given in Sections S2 B of the Supplementary Material. We find

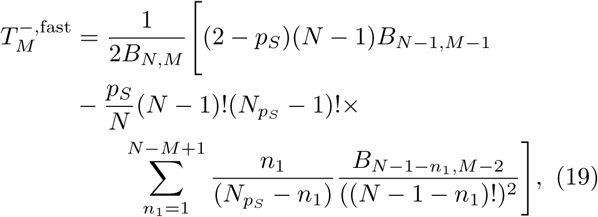

and

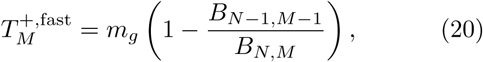

where the *B*_*k,l*_ = *B*_*k,l*_(*y*_1_, …, *y*_*k−l*+1_) are incomplete Bell polynomials as before. However, the *y*_*i*_ are now given by *y*_*i*_ = (*i −* 1)!(*N*_*ps*_*−* 1)_*i−*1_, where *N*_*ps*_ is

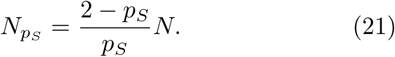

The stationary distribution for *M* is then obtained using Eqs. (13) and (14), with the replacement 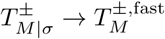.

We test these theoretical predictions in Figure 5 (c) and (d), and find good agreement with simulations. The behaviour of the model is further explored in Figure 6 (d-f), where we show the stationary distribution for the number of mating types in the limit of fast environments, for varying values of the facultative sex rate and for different mutation rates. For the parameters in Figure 6 the distributions are not too dissimilar from the ones in the slow switching regime (panels (a-c)). However, one main difference is the absence of bi-modality when *p*_*S*_ *≈* 1*/*2. Additionally, we find the distribution is wider around its peak.

### C. Transition between slow and fast switching regimes

In the previous sections, we studied the stationary distribution of the number of mating types in the limits of slow and fast environmental dynamics. The differences between these limits are most pronounced at intermediate facultative sex rates (*p*_*S*_ *≈* 1*/*2). We now focus on the regime of intermediate environmental switching. The time scale of the environmental dynamics is set by the cycle time *τ*, defined in Eq. (7), and we are therefore interested in situations where *τ* is comparable to the time scales of the evolutionary process in the population.

#### 1. Stochastic simulations

Results from simulations are shown in Figure 7. We focus on the case *p*_*S*_ = 1*/*2. Cases with *p*_*S*_ close to zero and one are explored in Section S3 C of the Supplementary Material. As shown in Figure 7, in the intermediate switching regime the stationary distribution 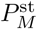 does not exhibit a clearly defined peak as in the fast switching regime, but rather it exhibits a wider distribution, with two peaks in some cases. When the population size is low (see first row of Figure 7), the distribution gets wider compared to the limiting cases (slow and fast switching regimes), and exhibits two peaks only when the mutation rate is low (see Figure 7 (a) for high values of *τ* ; this is the same distribution shown in Figure 4 (b)). For higher population sizes (see second row), the distribution makes a transition from a unimodal shape (fast switching regime) to bimodal (slow switching regime) for all values of the mutation rate used in the figure. In between (intermediate switching regime) the distribution is wider around its peak until it bifurcates in two. This situation is also observed for higher population sizes (see lower row), however, each of the two peaks become more narrow.

**FIG. 7.**
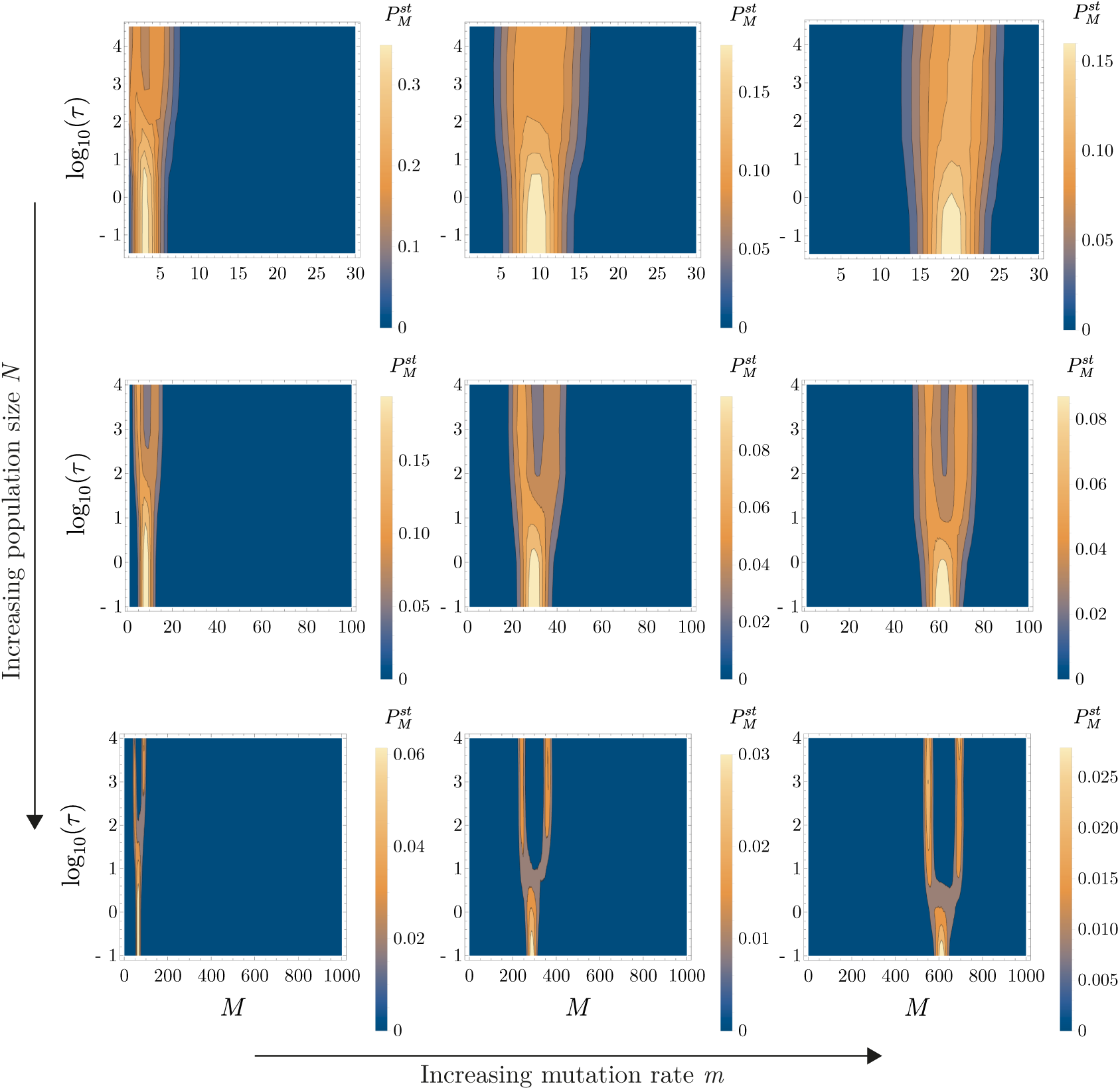
Simulation results of the stationary distribution 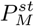 as function of *M* and *τ*, the average time of one switching period. Parameters used: upper row, *N* = 30; middle row, *N* = 100; lower row, *N* = 1000. Left column, *m* = 0.01; middle column *m* = 0.1; right column, *m* = 0.5. We set *λ*_*S→A*_ = *λ*_*A→S*_ throughout. Numerical simulations were conducted by time-averaging a long run until time *t* = 10^7^, with a time *t* = 10^6^ left to equilibrate.

#### 2. Generator-matrix approach

We have shown that in the single environment case, we are able to successfully reduce the combined dynamics of the number of mating types to an approximate one-step birth-death process for the number of mating types, *M* (see Section II C). We have further shown how this can be used to predict the population behaviour in the slow-switching limit (where the system takes on the average behaviour of the two independent environments, see Section III A), and in the fast-switching limit (where the system behaves as if it were in a single, effective environment, see Section III B). We now seek to extend this approach to the intermediate regime.

We begin by supposing that the dynamics of the full model (which involves transitions in the mating type abundances, **n**) can be approximated as a coupled birthdeath process in *M* and *σ*. The master equation for this process takes the general form

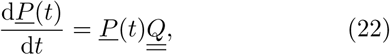

where the entries of the row vector *<u>P</u>* (*t*) are the probabilities of finding the system at a certain state (*M, σ*) at time *t*. It is convenient to arrange the states such that this vector takes the form

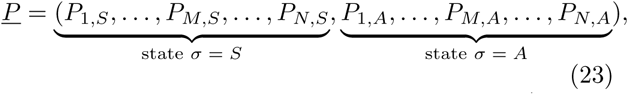

so the first half of entries correspond to states (*M, σ* = *S*), and the second one to states (*M, σ* = *A*). In both environments we have 1 *≤ M ≤ N*, with the bounds corresponding to the extreme cases of the whole population being of the same type (*M* = 1), or each individual of a different type (*M* = *N*). The matrix 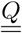 is of size 2*N ×* 2*N*, and it is convenient to write it in the following block structure,

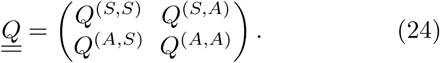

The *N × N* blocks *Q*^(*S,A*)^ and *Q*^(*A,S*)^ describe transitions between the environmental states. Given that the model does not include events in which both the environmental state and the number of mating types changes at the same time, these blocks are diagonal in *M*. We have 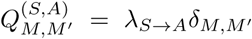 and similarly 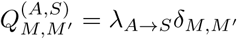. We must now find an approximation for the transitions within each environment, *Q*^(*S,S*)^ and *Q*^(*A,A*)^. Ultimately there are no correct choices for these matrices as they are simply approximations of the full model. Below we follow one particular approach, while alternatives are discussed in Section III C 3.

We begin by assuming that on transitioning to the sexual environment, the system rapidly relaxes to quasistationary state in which the dynamics of mating-type number are well-approximated by the transition rates in the sexual environment at equilibrium (see Section II C). Thus *Q*^(*S,S*)^ is tri-diagonal (only involving transitions that increase or decrease the number of mating types by one) with entries

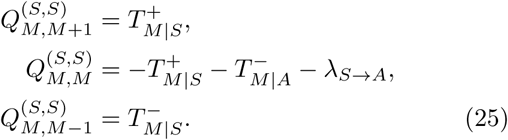

We can proceed analogously to construct the matrix *Q*^(*A,A*)^ in the asexual environment.

Next, we calculate the stationary distribution 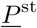 for this approximate system, by setting the right hand side of Eq. (22) to zero,

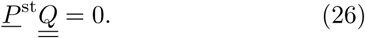

The solution has to be normalised appropriately, i.e, we must impose 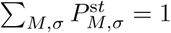. The marginal distribution for the number of mating types is obtained as 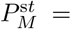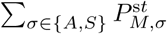. The block structure of 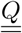 reduces the complexity of the problem, and, as a consequence, the stationary state can be obtained numerically relatively easily.

Compared to numerical simulations, the theoretical approach presented here brings a considerable simplification for estimating the stationary distribution 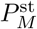. For the cases shown in Figure 7, simulations quickly become costly as both *N* and the switching rates increase, because more events occur per unit time. This is not a major obstacle, however, when solving Eq. (26). The matrix structure of 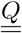 allows a fast numerical computation of 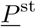 even for large values of *N*. In cases in which the distributions 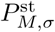 fall off quickly with *M* one can truncate the range of *M* to values much smaller than *N*, additionally accelerating the analysis.

We now proceed to define more precisely when exactly we expect this approach to work. We know that the key assumption is that the system has sufficient time in each environment to relax to the quasi-stationary distribution obtained in that environment when switching is absent. This assumption is required so that that we can approximate the rates 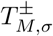 by 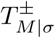 (a comparison to the case in which numerically-determined rates 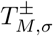are used in the generator matrix is presented in Section S2 E of the Supplementary Material).

In the sexual environment, this assumption is valid across a large parameter range (strong selection for even mating type frequencies very rapidly brings the system to a quasi-stationary distribution around one of the system’s fixed points). However in the asexual environment, relaxation to the stationary distribution takes far longer (this relaxation is driven entirely by genetic drift, which operates on a much slower timescale). Thus the requirement that this relaxation time is less than the typical time spent in the asexual environment provides the key restriction for the parameter range over which we expect the generator-matrix approximation to work. Assuming that the system in the sexual environment reaches 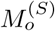 mating types before transitioning to the asexual environment, we now calculate the mean time taken for the system to relax from 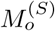 to 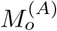 mating types in the asexual environment, where 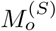 and 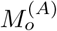 are the mode number of mating types in the fixed sexual and asexual environments, respectively. Using the results of [40] for a neutral multi-allelic Moran model, we find that the condition that the system spends sufficient time in the asexual environment is given by

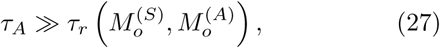

where

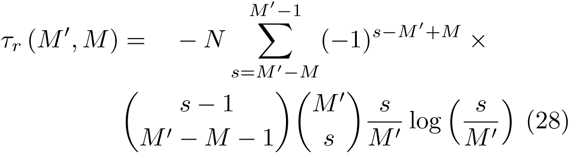

is the mean time taken for the system to transition from a state with *M*^*′*^ to *M* mating types in the asexual environment. Therefore, if the condition (27) is fulfilled, we expect that the system will have sufficient time to relax in the asexual environment to its quasi-stationary distribution for which 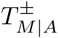 are accurate approximations for 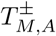.

In Figure 8, we show the predictions obtained using this approach for the same parameters as in Figure 7 (middle row). The parameters used are within in the range in which the generator-matrix approach is in good agreement with numerical simulations (see Eq. (27)). We note that this does not require very slow environmental switching per se; if the modes 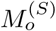 and 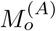 are sufficiently close, then intermediate switching rates allow the system to relax to the quasi-stationary distribution in the asexual environment. In fact, for the parameters used in Figure 8 the mean time *τ*_*r*_ is about ten times lower than *τ*_*A*_ for intermediate switching regimes. Thus if *m*_*g*_ is large (such that 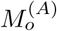 is large) or *N* low (such that 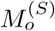 is low) we can still expect to see a good agreement between the generator-matrix approach and theory (see Figure 9). In biological terms we can therefore view the generator-matrix approach as being most useful when considering small populations, with migration (large *m*_*g*_) taking place between spatially segregated patches.

**FIG. 8.**
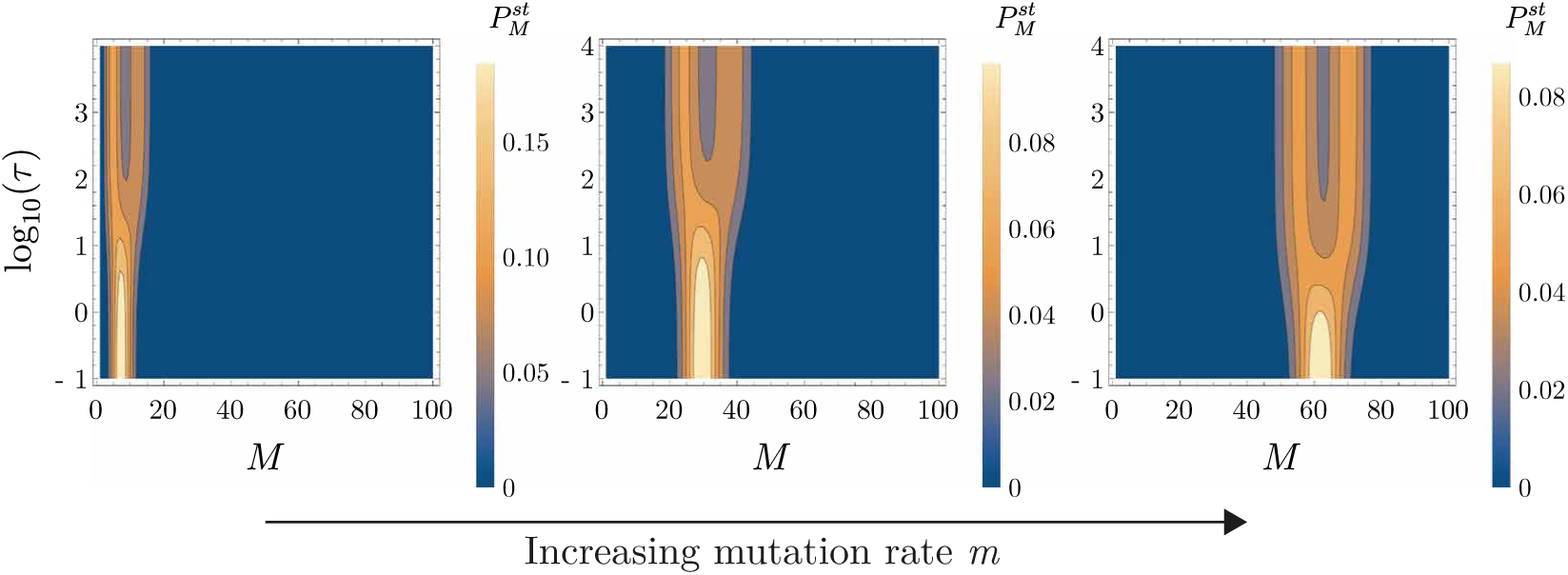
Theoretical prediction of 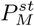 obtained as the solution of Eq. (26), i.e., the null space of matrix *Q*^*T*^. Parameters used are the same as in the second row of Figure 7.

**FIG. 9.**
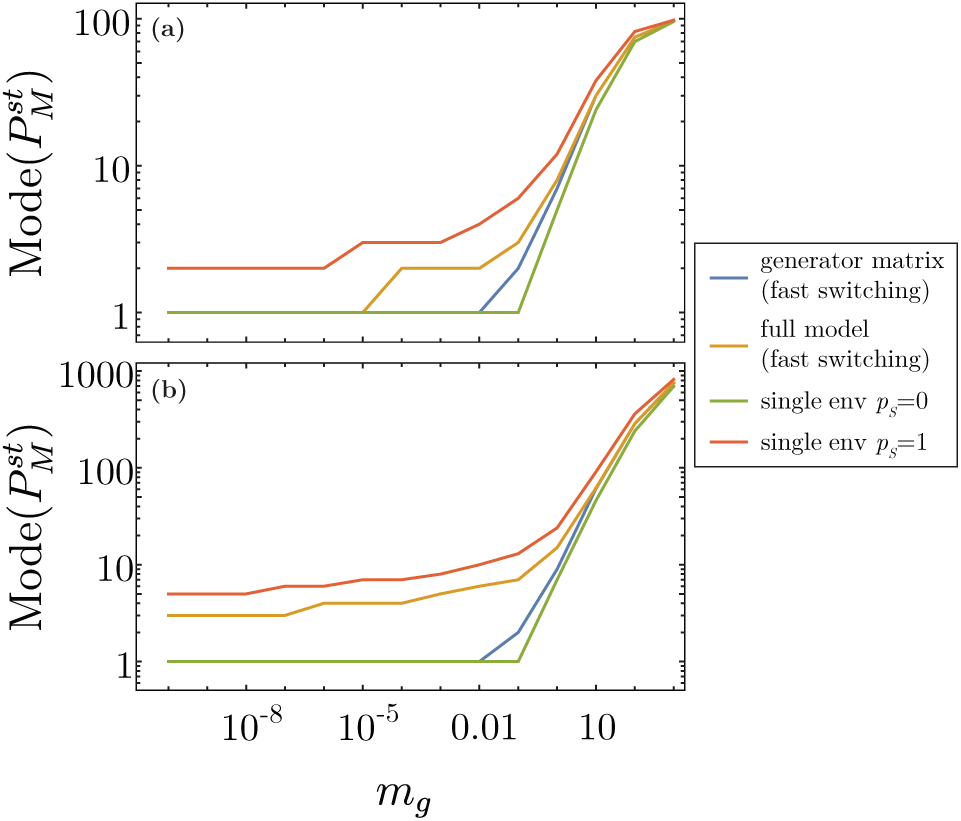
Mode of stationary distribution of the number of mating types, 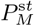, as function of *m*_*g*_ for *p*_*S*_ = 0.5 and population size: (a) *N* = 100 and (b) *N* = 1000. The prediction of the fast-switching limit in the full model is the approximation in Section III B, whilst the prediction of the generatormatrix approach in the fast-switching limit is described in Section S3 D of the Supplementary Material. The predictions of single environments with *p*_*S*_ = 0 and *p*_*S*_ = 1 correspond to the non-switching cases in Section S2.

In Section S3 A of the Supplementary Material we show in more detail how the distribution 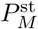 obtained from the generator-matrix approach compares against numerical simulations for different values of *m*_*g*_ and *p*_*S*_. We illustrate in Section S3 B how 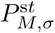 obtained from this approach behaves as function of *p*_*S*_ for both *σ* = *S* and *σ* = *A*. We also show that one can derive closed-form solutions of the stationary distributions 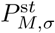 rates 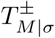 in the limits of slow and fast environmental dynamics (Section S3 D). In the slow-switching limit we obtain the result presented in Eq. (16). In this limit the system spends a long time in each environmental state, and thus, using the rates 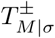 in the reduced model is a good approximation. For the fast-switching limit, however, this approximation is no longer valid; the system has insufficient time to relax to the quasi-stationary distribution in the asexual environment under which 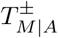 are accurate approximations for the transition rates, and so the theoretical prediction of 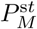 differs from the result presented in Section III B. Thus the generator-matrix approach can be understood as providing an approximation for the regime of slow-to-intermediate switching that improves on the slow-switching limit in Section III A.

#### 3. Alternatives to the generator-matrix approach

We have seen in Section III C 2 that while we can obtain a good approximation for the dynamics in the sexual environment in the environmental switching model, the approximation of the dynamics in the asexual environment is more challenging. This is perhaps surprising as in the asexual environment the dynamics of mating type frequencies are essentially given by a multi-allelic neutral Moran model, for which a wealth of well-established analytic results are available [40]. In this section we discuss two alternatives to the generator-matrix approach that leverage these results and demonstrate how, although initially plausible, each leads to their own set of issues.

First, we consider utilising standard results for the mean extinction time of a neutral allele. Assuming that on leaving the sexual environment with *M*^*′*^ mating types the frequency of each mating type is evenly distributed as *n*_*i*_ ≈ *N/M*^*′*^ (valid when *N* is large), the mean time to transition from *M*^*′*^ to *M* mating types is given by Eq. (28) in the absence of mutation. Given an initial number of mating types *M*^*′*^, the mean time to subsequently transition from *M* to *M −* 1 is then given by *τ*_*r*_(*M*^*′*^, *M −* 1) *−τ_r_*(*M*^*′*^, *M*). While this expression clearly features a dependence on the initial number of mating types in the asexual environment, *M*^*′*^, we find that this dependence is weak and in fact drops out in the limit of large *M*^*′*^. Assuming then that *m*_*g*_ is small, such that the probability that the probability of transitioning from *M* to *M* + 1 in the asexual environment is negligible, we can approximate the birth-death transitions in the asexual environment as

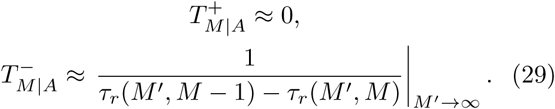

Here we have approximated the effective extinction rate of mating types, 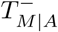, by inverse of the mean time to transition from *M* to *M −* 1.

While this may at first seem entirely reasonable, we find in fact that this model largely underestimates the number of mating types seen in the full model. The central problem is that while the mean transition time implied by Eq. (29) does indeed approximate the mean transition time in the full model, the full distribution of transition times is poorly predicted. Equation (29) assumes that the waiting time for a transition from *M* to *M −* 1 is exponentially distributed, with a non-zero probability of transitioning after a very small time in the asexual environment. However, the real distribution of transition times is peaked at a particular time, with transitions at very small times being impossible (it takes a minimum of *N/M^′^* reproductive events to drive a mating type extinct). In this way the approach suggested in Eq. (29) allows more frequent extinctions than actually observed, and thus a lower number of mating types in the stationary distribution than we see in simulations.

A second approach is to ignore the distinct asexual environment entirely, but to instead allow arbitrary transitions from a state *M*^*′*^ to all states *M < M^′^*. Again, we assume that *m*_*g*_ is small, and ignore the possibility of an increase in the number of mating types in the asexual environment. In the sexual environment, we have contributions to the probability that the number of mating types increases or decreases by one, as in Section III C 2. When the system enters the asexual environment, we now ask what is the probability of transitioning from *M*^*′*^ *M* before the system reverts to the sexual environment. In this way we can circumvent any direct modelling of the asexual environment while slightly increasing the complexity of the single-environment model by adding nonlocal transitions.

While the above approach may at first seem analytically challenging, progress is in fact possible. In [40], an expression was developed for the probability that exactly *M* alleles remain in a population at some time *t*. All we need to do is integrate this function over the probability of transitioning from the asexual to the sexual environment at time *t* (i.e., the exponential distribution with parameter *λ*_*A→S*_). We find then that accounting for the asexual environment yields the following contribution to the probability per unit time of transitioning from *M* to *M*^*′*^:

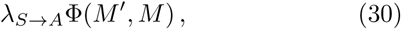

with

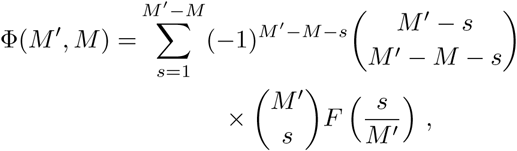

where

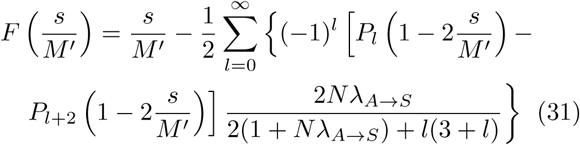

and *P*_*l*_ (*y*) are Legendre polynomials.

The above technique provides an analytically elegant alternative to the generator-matrix approach. Unfortunately, it turns out to be numerically impractical. Equation (31) involves the infinite sum over Legendre polynomials, and the slow convergence of these terms is a known numerical issue [41]. Convergence is especially problematic when *Nλ_A→S_* is large, a range (large population size) that is particularly interesting biologically.

Therefore while this second approach has the best potential for providing an analytic approximation to the number of mating types at intermediate regimes, its ultimate success relies on an improved analytic or numerical method for tackling Eq. (31), which lies outside the scope of this paper.

#### 4. Regime of small m_g_ and large N

Having developed approximations for the intermediate regime when *m*_*g*_ is large and *N* is small, we here investigate the range and extent of the intermediate switching regime when *m*_*g*_ is small and *N* is large, reflecting a more panmictic population in which *m*_*g*_ can readily be interpreted as a mutation rate. In Figure 10 we plot the mode number of mating types in the stationary distribution as a function of the probability of being in a sexual environment, *p*_*S*_, for varying mean residency times in the sexual state, *τ*_*S*_. We see that for *p*_*S*_ *≈* 1 (almost obligate sex) the number of mating types is well described by the fast-switching theory of Section III B. However as *p*_*S*_ is lowered, we begin to see departures from this theory, with the number of mating types consistently lower than that predicted by the fast-switching limit. These departures are ever more extreme as the time spent in the sexual environment (and consequently for fixed *p*_*S*_, also the time spent in the asexual environment) increases. We can therefore see that fast-switching theory very much represents an upper-bound on the mode number of mating types expected for a general set of parameters.

**FIG. 10.**
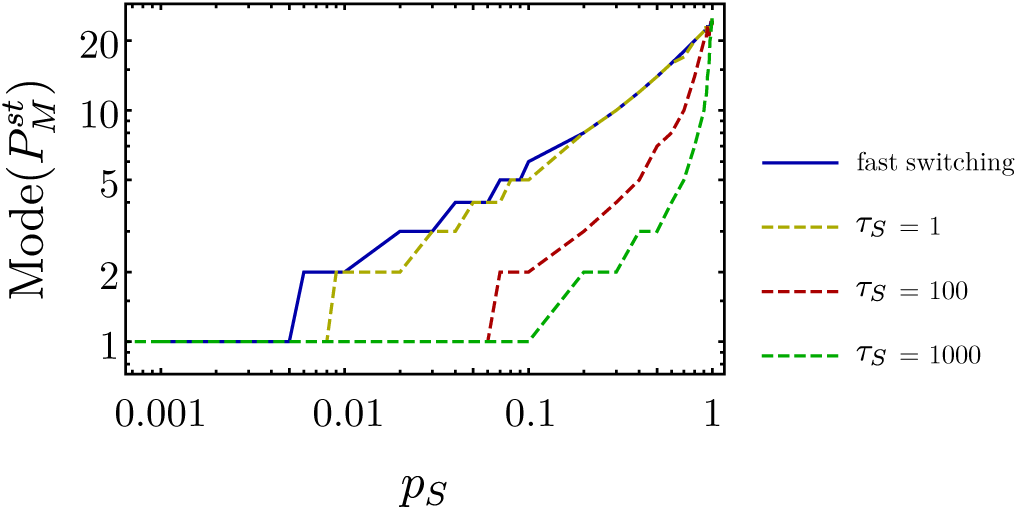
Mode of stationary distribution of the number of mating types, 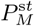, as function of the probability of being in a sexual environment, *p*_*S*_, for parameters *N* = 10^4^ and *m*_*g*_ = 10^−3^. The prediction of the fast-switching regime is based upon the results studied in Section III B. For longer residency times in the sexual environment (i.e., longer *τ*_*S*_), simulations of the full model (dashed lines) demonstrate a lower number of mating types, in qualitative agreement with the results of Section III C 2. Discrepancy between the fast-switching limit and the simulations increases as *p*_*S*_ increases, when the system enters the intermediate switching regime. As *p*_*S*_ approaches zero, the system enters the slow-intermediate regime described by the generator-matrix approach (see Eq. (27)), in which only one mating type can be maintained.

**FIG. 11.**
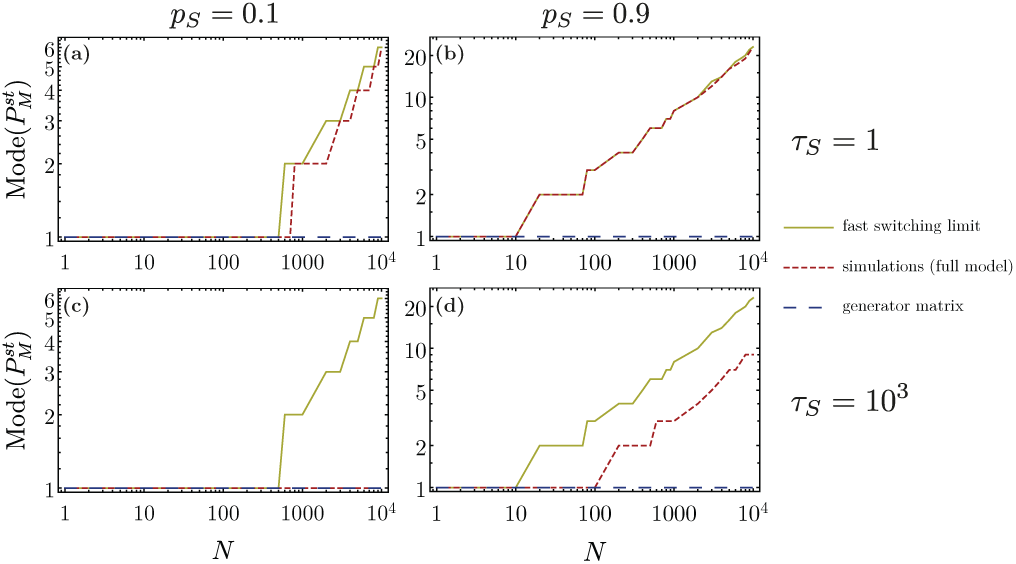
Mode of stationary distribution of the number of mating types, 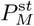, as function of the population size *N* for *m*_*g*_ = 10^−3^ for different values of *p*_*S*_ and *τ*_*S*_. The fast-switching prediction is based upon the results studied in Section III B, whilst the generatormatrix approach upon the pre-diction of the framework presented in Section III C 2. Simulations were run up to time *t* = 10^8^, with measurements starting *t* = 10^3^ to ensure stationarity.

While it is computationally impractical to investigate population sizes much larger than *N* = 10^4^, or mutation rates much lower than 10^−3^ (as in Figure 10), we are nevertheless interested in what general patterns we might expect to see as we go beyond this regime. In Figure 11 we investigate how the mode number of mating types varies with population size. Broadly our results fit our intuition developed thus far; when the time spent in the asexual environment is very short (see panel (b)) the system is well-approximated by the fast switching theory, while when the time spent in the asexual environment is very long (see panel (c)) the generator-matrix approach works well (in fact, *τ*_*r*_ in Eq. (28) is less than one percent of *τ*_*A*_ for all the values of *N* shown in panel (c)). Meanwhile at intermediate switching regimes we see that the magnitude of departure from the fast-switching theory increases as time spent in the asexual environment increases (see panels (a) and (d)). However we also see now that as *N* increases, the simulation results for intermediate switching rates begin to approach the fast-switching limit. In the context of Eq. (28), this is perhaps unsurprising; the timescale on which extinctions occur in the asexual environment is linearly dependent on *N*, and so, as this population size increases, the range of switching rates that can still be considered fast also increases.

## IV. SELECTIVE SWEEPS IN THE SWITCHING ENVIRONMENTAL MODEL

In the previous section we demonstrated that explicitly accounting for switching environments led to both quantitative and qualitative changes in the model predictions. In this section we will show how this change in modelling formalism allows us to tackle a richer array of biological questions, without necessarily sacrificing tractability.

Suppose that mutations arise in the population at loci unlinked to the mating type locus at an average rate *µ*. We will further suppose that the mutant allele is under directional (frequency independent) selection, such that individuals carrying the mutation have a selective advantage *s* over individuals carrying the resident allele. If *s <* 0, the mutation will be selected against and will be rapidly lost from the population. If *s >* 0 the mutation will be selected for and (in the absence of stochastic extinction effects) will sweep to fixation. However the focus of interest for this study is the frequency of the mating type alleles. The impact of this selective sweep will have very different effects on the mating type frequencies depending on the environment, sexual or asexual, in which it occurs.

If this mutation arises while the system is in the sexual environment, it quickly spreads to all the present mating types via genetic recombination and has no effect on the number of mating types *M*. For instance, say that mutation occurs in an individual of mating type 1. While that individual experiences a selective advantage *s*, upon sexual reproduction with a nonself mating type (say of mating type 2) the beneficial mutation will have the opportunity to spread to another mating type class. Thus the beneficial mutation will rapidly spread through the mating type populations without appreciably distorting their frequencies.

Conversely, if this mutation arises while the system is in the asexual environment, it is confined to the mating type on which it occurs (genetic recombination is absent) and will cause large distortions in the mating type frequencies. Denoting by *y* the frequency of individuals carrying the beneficial mutation, their dynamics in the large *N* limit will be given by

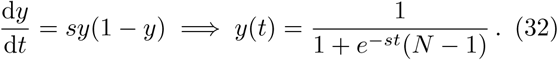

Let us suppose again that the beneficial mutation occurs in an individual of mating type 1. Then, assuming that initially the mating types were in approximately equal abundances, the dynamics for the mating type frequencies *x*_*i*_ is given by

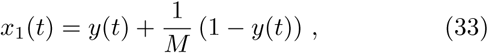

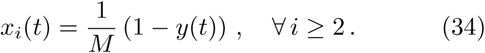

These dynamics are illustrated in Figure 12. We see that as the mutation sweeps through the population, the total number of individuals of mating types *i ≥* 2 decreases, and thus, stochastic extinctions of these mating type classes become more likely. Over only slightly longer timescales, fixation of a single mating type is all but guaranteed, with the mean time until the fixation of a single mutant given approximately by 2 log(*N*)*/s* in large populations. We now leverage the results of the previous section to provide a more simple approach using the reduced model. We explain this below.

**FIG. 12.**
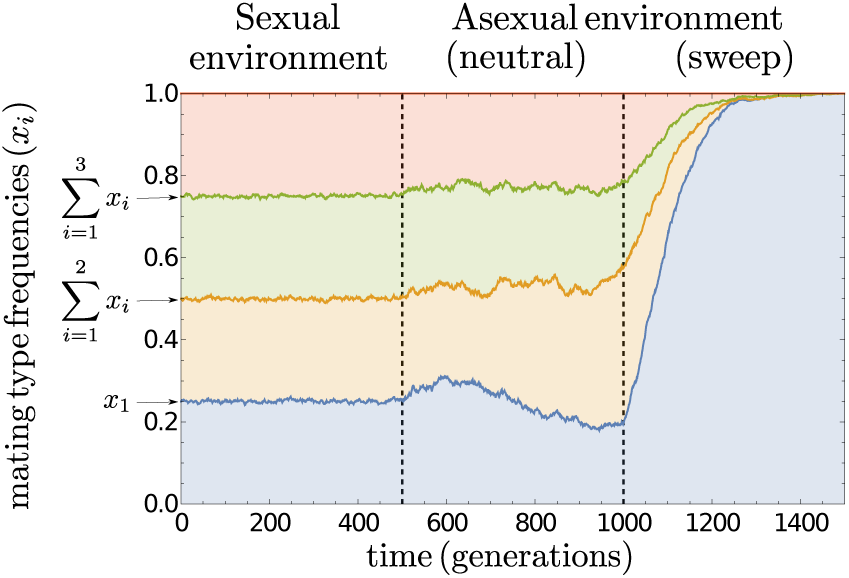
Illustration of the dynamics of mating type frequen-cies *x*_*i*_ = *n*_*i*_/*N* when selective sweeps are accounted for. In the sexual environment (0 *< t <* 500) mating type frequen-cies are held at approximately equal values by negative fre-quency dependent selection. Beneficial mutations at unlinked loci spread to each mating type subpopulation through re-combination, and thus do not affect this even mating type distribution. In the asexual environment (500 *< t <* 1500), in the absence of beneficial mutations (500 *< t <* 1000) mating type frequencies fluctuate due to genetic drift alone. However when beneficial mutations occur (*t* = 1000), the mating type background on which they arise can hitchhike to fixation, re-ducing the number of mating types to one. Data is obtained from Gillespie simulation with *N* = 5 ×10^4^ and *s* = 0.02.

In addition to the effective birth-death and switching environment processes previously present in the generator-matrix approach, we now consider the possibility that the system transitions to a state of one single mating type at constant rate *ν* when in the asexual environment. The rate *ν* can thus be understood as a compound parameter that captures the average rate at which a beneficial mutation occurs in an asexual environment (*µp_S_*) and has sufficient time to sweep a single mating type to fixation. this occurs with probability 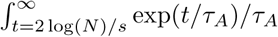, and we therefore have

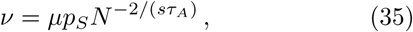

where 2 log(*N*)*/s* is the conditional mean fixation time for a single mutant to fixate in large populations. We consider the rates 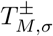 as before, first assuming a fixed environment. For environment *σ* = *A* then, we have

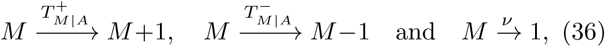

while for *σ* = *S*

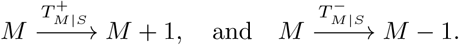

The switching environment transitions are as in the previous sections (i.e., as in Eq. (1)). The stationary distribution 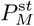 for this scenario can be estimated in a similar way to the method presented in Section III C 2, by constructing the corresponding generator matrix 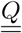 in which the selective sweeps process 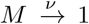 is included (see Section S4 A of the Supplementary Material for details). Figure 13 shows the theoretical prediction of 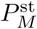 obtained from this approach for both switching environments, considering an equal fraction of time spent in each environment (i.e., *p*_*S*_ = 1*/*2).

**FIG. 13.**
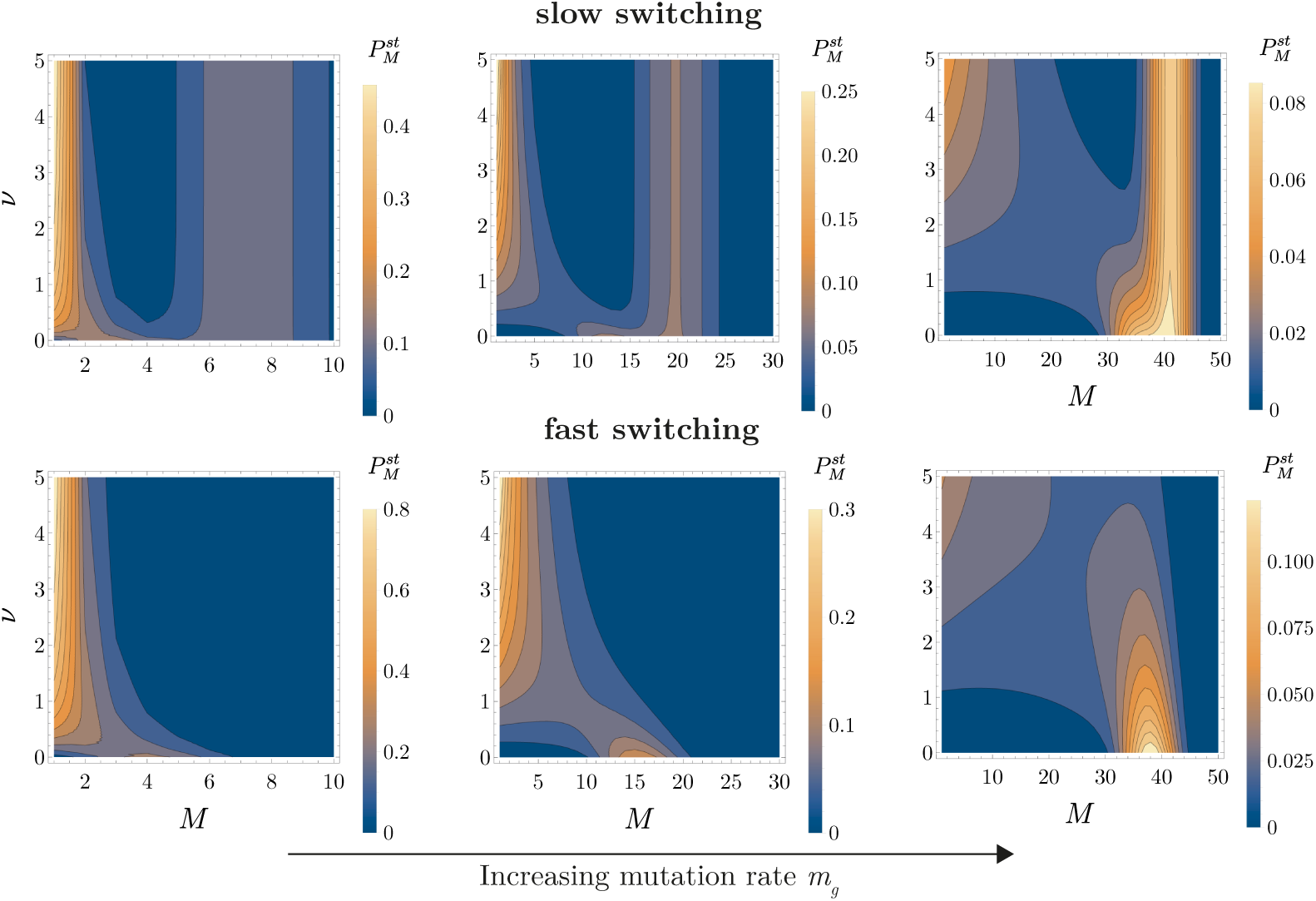
Theoretical prediction of 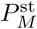 as function of *M* and *ν* for different values of *m* in slow and fast switching environments. Population size *N* = 50. From left to right panels: *m*_*g*_ = 0.5, 5, 5.

The inclusion of selective sweeps brings interesting features in both switching regimes. For the slow-switching case (see upper row of Figure 13), we observe that the distribution still remains bimodal but with a considerably higher peak at *M* = 1. As *ν* increases, the distribution at both modes remains constant. On the other hand, for the fast-switching regime (see lower row of Figure 13), as *ν* increases the mode transitions from a value *M >* 1 to *M* = 1. In both regimes, the emergence of the peak at *M* = 1 occurs when *ν* crosses certain point determined by how high the mutation rate *m*_*g*_ is. As *m*_*g*_ increases, this point will naturally be higher. The approach employed here to predict 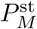 is an approximation as it makes use of rates 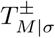 that assume a fixed environment. We compare it against numerical simulations in Section S4 A in the Supplementary Material. We also explore the case of selective sweeps in a fixed asexual environment in Section S4 B. Our theoretical predictions capture the main effects on the distribution of the number of mating types when selective sweeps are included.

In the presence of selective sweeps we can then see the following biological picture emerge. In order for the populations to maintain more than a single (essentially non-functional) mating type, one of two scenarios must hold. In the first scenario we see *m*_*g*_ *» ν*. In this case the rate of supply of new mating types (governed by *m*_*g*_) far exceeds the extinction rate generated by selective sweeps (governed by *ν*). This would be appropriate if we were to consider *m*_*g*_ as representing a migration rate between geographically structured subpopulations. In the second scenario we see *s*^*−1*^ *> τ*_*A*_. In this case while selective sweeps can initiate in the asexual environment, switching is sufficiently fast that the sweeps cannot complete.

## V. CONCLUSIONS

For reasons on tractability, most studies considering evolution in facultatively sexual populations focus on asynchronous sex, in which individuals probabilistically engage in sexual reproduction [29]. This is also true for models that have tried to capture the evolution of mating type number under demographic stochasticity [24, 25, 30]. In this paper we have released this restriction to consider the dynamics of the number of mating types under demographic stochasticity in populations that switch synchronously between asexual and sexual environments. In a course grained sense our model recapitulates previous theoretical and empirical observations that the number of mating types should be positively correlated with increasing amounts of sexual relative to asexual reproduction. However we have shown that the additional consideration of sexual synchrony generates both quantitative and qualitative differences from the asynchronous model, as well as offering scope for asking a richer array of biologically interesting questions.

With respect to quantitative differences between the asynchronous and synchronous models, we have shown the two models are only equivalent in the limit of fast switching between environments. However as switching becomes slower (and in particular as the amount of time in the asexual environment becomes longer) mating type extinctions become more likely in the synchronous model, lowering the expected number of mating types in the stationary distribution. For instance, in Figure 10 with a probability of *p*_*S*_ = 1*/*2 of being in the sexual environment, we see a reduction of more than 10 mating types when the time spent in the sexual environment is large (*τ*_*S*_ = 100) relative to the asynchronous (fast-switching) theory. This reduction may explain previous overestimates in the expected number of mating types when compared with previous studies where asynchronous sexual reproduction was assumed [24, 25]. In fact the mode number of mating types can drop to just one type over a range of biologically relevant parameters.

Qualitative differences between the asynchronous and synchronous models are most apparent in small populations. In this scenario the parameter *m*_*g*_ can be interpreted as a per-generation migration rate (with mating types coming from a highly diverse pool), which we expect biologically to be much higher than a mutation rate. When switching is fast, we see as before that the model tends to the limit of asynchronous switching. However, when switching is slow, a bimodal prediction for the number of mating types is possible. Here the population spends enough time in the asexual and sexual environments that the stationary distribution approaches a superposition of those in the fixed environments; just a single mating type is maintained in the sexual environment, while in the sexual environment ingressing mating types rapidly establish. We emphasise that this behaviour is not possible in the asynchronous model. While local absences of particular mating types are common in samples of fungi (e.g. *Coccidioides* [42]) and ciliates (e.g. *Tetrahymena pyriformis* [43]), obtaining empirical distributions of the number of mating types across geographic locations is hindered by low sample sizes. However our analysis agrees qualitatively with the observation that the presence of more than one mating type is indicative of more recent sexual activity [44]. Meanwhile observations of all mating types present across geographic regions (as for instance in *Dictyostelium discoideum* [45]) is consistent not only with the observation of relatively high rates of sexual reproduction, but also of a fast switching rate between asexual and sexual reproductive modes.

From a modelling perspective, by allowing for synchronous sexual reproduction we have also been able to tackle the issue of selective sweeps. In [30] it was shown that selective differences between mating types (induced, for instance, by non-neutral mutations at loci linked to the mating type locus) could rapidly reduce the number of types observed. As the model was deterministic and assumed asynchronous sexual reproduction, it could not quantify how selective sweeps (caused by beneficial mutations at loci unlinked to the mating type locus) might affect the the number of types observed in isogamous species. Experimentally however, such sweeps have been shown to be strong drivers of mating type extinctions in facultatively sexual species such as *Chlamydomonas* [31] and *Tetrahymena* [46]. Accounting for this effect mathematically, we have been able to show that although this effect decreases rapidly with increasing population size, a substantial extinction risk is present when the product of the strength of beneficial mutations and the average time spent in the asexual environment is greater than one (*sτ_A_ >* 1).

By accounting for demographic stochasticity, synchronous sex, and selective sweeps, our model suggests that the persistence of self-incompatible mating types in facultatively sexual populations may be even more precarious than previously believed [39]. In fact, a range of biologically plausible parameters suggest that just a single (functionally asexual) mating type is most probable at long times. Empirical phylogenies of isogamous species such as within fungi [47] and ciliates [48] show that such scenarios are relatively common. And yet despite this, species with distinct mating types have remained stable over long evolutionary periods, with highly conserved mating type loci [49]. In light of this it is possible to reframe the evolutionary question away from asking “*Why do most isogamous species have just two mating types?*” and towards “*How do so many facultatively sexual species maintain even two in the face of genetic drift and selective sweeps?* “. Our bimodal results for small population sizes with an effective migration rate point to a possible solution, suggesting a role for spatial structure. If sex is not synchronised in time across a whole population, but rather synchronised across a finite set of spatial regions, mating type diversity may be maintained. However, further work, perhaps involving metapopulation models [50], would be needed to fully uncover how geographic population structure might affect the evolution of mating type number.

More broadly, our results also point to some important considerations in the general literature on the evolution of sexual reproduction. While the facultative nature of sexual reproduction in models that assume asynchronous sex is captured by just a single parameter (the probability of a sexual rather than asexual event), models of synchronous sex require the specification of two parameters (the mean time spent in sexual and asexual environments). For species in which sexual reproductive phases are induced in a seasonal or regular manner (such as *Tetrahymena* [51]), these parameters may be relatively straightforward to estimate. However for species in which sexual reproduction is irregular or very rare, such as (*Chlamydomonas* [52] or *Saccharomyces* [53]) estimating these parameters independently may pose a far greater challenge. Here ‘rates of sexual reproduction’ are often estimated using genomic methods that can be used to infer the long-time average number of sexual to asexual reproductive events (equivalent to *p*_*S*_ in our model), but the of duration time spent in each state is left unspecified [53, 54]. While difficult to obtain, we suggest that obtaining estimates of these parameters is a worthwhile endeavour, as we have shown specifying their precise values can have important evolutionary consequences.

One example, with a direct relation to the current study, is the evolution of ‘mating type switching’, the ability of individuals to change their expressed mating type between asexual reproductive events. This has been previously explored using simulations of populations with synchronous sexual reproduction [23], with the number of asexual generations between single sexual generations varied. It was found that mating-type switching was more likely to evolve as the number of concurrent asexual generations increased (which distorted the relative frequencies of non-switching mating types through drift, and placed them at a selective disadvantage). While we also see strong distortions in mating type frequencies here when periods of asexual reproduction are long (large *τ*_*A*_), it should be noted that we also observe distorted frequencies when the probability of being in the sexual environment is very small (*p*_*S*_ = *τ*_*S*_/(*τ*_*S*_ + *τ*_*A*_)). Thus, long periods of asexual reproduction may not be needed for the evolution of mating type switching.

A second example is the enigma of the evolution of sexual reproduction and genetic recombination itself [55]. It has been suggested that facultative sex provides the best of both worlds [56], engendering species with the benefits of sexual reproduction (increased genetic diversity and evolvability [57]) while minimising the costs (for instance finding a suitable mate or the ubiquitous the costs of recombination [58]). In this sense it has been further suggested that in maximising the possibilities of finding a mate, synchronous sexual reproduction may be better still [29]. However, as we have shown, asynchronous sex can lead to its own costs at the population level in terms of an increased extinction probability of partners with which to mate. Thus, in the absence of a mechanism to provide assured sexual reproductive opportunities in later generations (such as mating type switching), some level of asynchronous sex may in fact be beneficial for maintaining the diversity of compatible partners at the population level.

In evolutionary modelling there is always a natural tension between analytic tractability and biological realism. The optimal point between these two extremes is ultimately subjective, however, an increased level of biological realism is arguably warranted if it: (i) generates qualitative differences; (ii) generates quantitative differences or (iii) allows the exploration of more interesting biological questions. Under these metrics, we have shown that accounting for the synchrony of sex is an important modelling consideration for investigating the evolutionary dynamics of the number of mating types in finite populations. We have demonstrated that while analytic results can be derived in a relatively straightforward manner in the fast and slow switching regimes, intermediate switching rates pose more of an analytic challenge. While improvements on the slow-switching approximation through the generator-matrix approach are possible under certain conditions, a more generally applicable approximation is difficult to obtain. This is, as we have discussed, despite the relative simplicity of the neutral dynamics in the asexual environment. Developing further analytic methods for dealing with systems featuring both demographic stochasticity and switching environments presents an interesting mathematical challenge, but also one that is ultimately required for a full understanding evolution in facultatively sexual species. It is our hope that in the coming years this challenge will be taken up to yield new evolutionary insight on a host of problems involving facultative sex.

## ACKNOWLEDGEMENTS

EBC acknowledges a President’s Doctoral Scholarship (The University of Manchester). TG is grateful for partial financial support by the Maria de Maeztu Program for Units of Excellence in R&D (MDM-2017-0711). GWAC thanks the Leverhulme Trust for funding through a Leverhulme Early Career Fellowship. The authors thank the International Centre for Mathematical Sciences (ICMS, Edinburgh, UK). This work was initiated at their workshop ‘Stochastic models of evolving populations: from bacteria to cancer’ (July, 2018).

## Notes

### Competing Interest Statement

The authors have declared no competing interest.

