## Supplementary Material for "Switching environments, synchronous sex, and the evolution of mating types"

### Switching environments, selective sweeps and the evolution of the number of sexes
### — Supplementary Material —

**Ernesto Berríos-Caro, Tobias Galla, and George W. A. Constable**

#### CONTENTS


S1. Effective rates $T^{-}_{M|\sigma}$ and $T^{+}_{M|\sigma}$ as function of $P^{\text{st}}_{\mathbf{n}|\sigma}$ S2
- A. Calculation of the effective rate $T^{-}_{M|p_S}$ S3
- B. Calculation of the effective rate $T^{+}_{M|p_S}$ S5

S2. Closed-form solutions of $T^{-}_{M|p_S}$, $T^{+}_{M|p_S}$, and $P^{\text{st}}_{M|p_S}$ S5
- A. Asexual environment ($p_S = 0$) S6
  1. Calculation of $S_1 = \sum_{\mathbf{n}\in\mathcal{S}^M} P^{\text{st}}_{\mathbf{n}|A}$ S7
  2. Calculation of $S_2 = \sum_{\mathbf{n}\in\mathcal{S}^M} \left(\sum_{i=1}^{M} \delta_{n_i,1}\right) P^{\text{st}}_{\mathbf{n}|A}$ S7
  3. Rates $T^{-}_{M|A}$ and $T^{+}_{M|A}$, and stationary distribution $P^{\text{st}}_{M|A}$ S8
- B. Facultative sex ($0 < p_S < 1$) and sexual environment ($p_S = 1$) S9
  1. Calculation of $S_1 = \sum_{\mathbf{n}\in\mathcal{S}^M} P^{\text{st}}_{\mathbf{n}|p_S}$ S9
  2. Calculation of $S_2 = \sum_{\mathbf{n}\in\mathcal{S}^M} \left(\sum_{i=1}^{M} \delta_{n_i,1}\right) P^{\text{st}}_{\mathbf{n}|p_S}$ S11
  3. Calculation of $S_3 = \sum_{\mathbf{n}\in\mathcal{S}^M} \left(\sum_{i=1}^{M} n_i^2\right) \left(\sum_{i=1}^{M} \delta_{n_i,1}\right) P^{\text{st}}_{\mathbf{n}|p_S}$ S11
  4. Rates $T^{-}_{M|p_S}$ and $T^{+}_{M|p_S}$, and stationary distribution $P^{\text{st}}_{M|p_S}$ S13
- C. Alternative expressions of rates $T^{-}_{M|p_S}$ and $T^{+}_{M|p_S}$ for cases $0 < p_S \leq 1$ S13
- D. Comparison against numerical simulations with fixed environment S16
- E. Comparison against numerical simulations with switching environments S16

S3. Markov chain generator matrix approach S18
- A. Comparison against simulations S18
- B. Theoretical prediction of $P^{\text{st}}_{M,\sigma}$ for each environment S19
- C. Transition between slow and fast switching regimes for low and high $p_S$ S19
- D. Analytical results from the generator-matrix approach S20
  1. Limit of slow environmental switching S21
  2. Limit of fast environmental switching S22

S4. Selective sweeps S24
- A. Switching environments S24
- B. Fixed asexual environment S26

References S27

#### S1. EFFECTIVE RATES $T^-_{M|\sigma}$ AND $T^+_{M|\sigma}$ AS FUNCTION OF $P^{\mathbf{st}}_{\mathbf{n}|\sigma}$

In this section, we provide details for the first step of the calculation of the rates $T^-_{M|\sigma}$ and $T^-_{M|\sigma}$ in the reduced model. These are defined in the main paper. Throughout the calculation we assume that the environmental state is fixed in time. This means that the birth and death rates in the full model do not vary with the environment. The result of this section is expressed in terms of the stationary distribution of states $\mathbf{n}$ of the full model, $P^{\text{st}}_{\mathbf{n}|\sigma}$. Further evaluation then follows in the subsequent section of this Supplementary Material.

To derive results for the effective rates in the reduced model we start from the transition rates from the full model. The rates for birth-death events are given by

$$\begin{aligned} \mathcal{T}^S_{ij} &= \frac{1}{2}\frac{n_i n_j}{N^2}\left(N - n_i\right), \\ \mathcal{T}^A_{ij} &= \frac{n_i n_j}{N} \end{aligned} \tag{S1}$$

in the two environmental states ($S$ and $A$). This describes rate of events in which one individual of mating type $i$ reproduces and an individual of type $j$ is removed, i.e., events in which $n_i \to n_i + 1$ and $n_j \to n_j - 1$.

Mutation events occur with rates

$$\mathcal{T}^m_j = m_g \, \frac{n_j}{N}, \tag{S2}$$

independent of the state of the environment.

We carry out the calculation at a more general level than needed for the system with two environmental states. Following [1], we allow for an arbitrary rate of sexual reproduction, $0 \le p_S \le 1$, and use

$$\mathcal{T}^{p_S}_{ij} = \left[(1-p_S)\frac{n_i}{N} + \frac{p_S}{2}\frac{n_i}{N}\left(\frac{N-n_i}{N}\right)\right] n_j, \tag{S3}$$

instead of the rates given in Eq. (S1). We note that time is measured in units of generations in our model, so that the transition rates $\mathcal{T}^{p_S}_{ij}$ carry an extra factor $N$ in comparison to [1].

The specific cases of fully sexual or fully asexual reproduction ($\sigma = S, A$) can be obtained by setting $p_S = 1$ or $p_S = 0$ respectively. We then recover the transition matrices defined in Eqs. (2) and (3) of the main paper as

$$\mathcal{T}^{p_S=1}_{ij} = \mathcal{T}^S_{ij} \quad \text{and} \quad \mathcal{T}^{p_S=0}_{ij} = \mathcal{T}^A_{ij}. \tag{S4}$$

We write $T^-_{M|p_S}$ and $T^+_{M|p_S}$ for the effective rates of the reduced model, when the full model is such that birth-death events occur with the rates given in Eq. (S1). Our aim is to calculate the $T^\pm_{M|p_S}$, when $p_S$ is fixed in time in the full model.

##### A. Calculation of the effective rate $T^-_{M|p_S}$

For small $\Delta t$, the quantity $\Delta t\, T^-_{M|p_S}$ can be obtained from the probability that the number of mating types decreases by one in the next $\Delta t$ units of time, given that there are currently $M$ mating types. More precisely,

$$P^-_{M|p_S}(\Delta t) \equiv P(\{M \to M-1\}|M \text{ mating types in the population }) = \Delta t\, T^-_{M|p_S} + \mathcal{O}(\Delta t^2). \quad \text{(S5)}$$

In this expression we have written $\{M \to M-1\}$ for the event in which the number of mating types reduces from $M$ to $M-1$ in the next $\Delta t$ units of time. We always assume a fixed value of $p_S$ in the full model.

When $M$ different mating types are present in the population, the system can be found in any state $\mathbf{n}$ that has $M$ non-zero entries. We write $C_\mathbf{n}$ for the event that the population is in state $\mathbf{n}$. Since the population size is fixed at $N$, the sum of the entries of $\mathbf{n}$ is always $N$. The vectors $\mathbf{n}$ of this type are known 'compositions' (or '$M-$compositions') of the integer $N$ [2]. We write $\mathcal{S}^M$ for the set of these compositions, i.e., $\mathcal{S}^M$ is the set of vectors $\mathbf{n}$ with precisely $M$ non-zero entries, and with $\sum_i n_i = N$.

To ease the notation we introduce the following events (in the sense of probability theory):

$$\begin{aligned} A &= \text{the event in which the number of mating types reduces from } M \text{ to } M-1 \\ &\quad\ \text{in the next } \Delta t \text{ units of time.} \\ B_M &= \text{the event to find the population in a state with } M \text{ mating types} \\ C_\mathbf{n} &= \text{the event to find the population in state } \mathbf{n}. \end{aligned} \quad \text{(S6)}$$

In this notation, $P^-_{M|p_S}(\Delta t) = P(A|B_M)$. We note that $B_M = \cup_{\mathbf{n}\in\mathcal{S}^M} C_\mathbf{n}$, and that the $C_\mathbf{n}$ are pairwise disjoint. The conditional probability in Eq. (S5) can then be written as

$$\begin{aligned} P^-_{M|p_S}(\Delta t) &= \sum_{\mathbf{n}\in\mathcal{S}^M} P\left(A \cap C_\mathbf{n} | B_M\right) \\ &= \sum_{\mathbf{n}\in\mathcal{S}^M} \frac{P\left(A\cap C_\mathbf{n} \cap B_M\right)}{P(B_M)}. \end{aligned} \quad \text{(S7)}$$

Noting that $C_\mathbf{n}$ is a subset of $B_M$ for $\mathbf{n} \in \mathcal{S}^M$, we have $P\left(A\cap C_\mathbf{n}\cap B_M\right) = P\left(A\cap C_\mathbf{n}\right) = P(A|C_\mathbf{n})P(C_\mathbf{n})$. Using this, we find

$$P^-_{M|p_S}(\Delta t) = \sum_{\mathbf{n}\in\mathcal{S}^M} P\left(A|C_\mathbf{n}\right) \frac{P(C_\mathbf{n})}{P(B_M)}. \quad \text{(S8)}$$

This formula has a straightforward interpretation. We are interested in the probability that the event $A$ occurs (extinction of a mating type in the next $\Delta t$), given that there are currently $M$ mating types in the population (condition $B_M$). If there are exactly $M$ mating types in the population, then the population is in state $\mathbf{n} \in \mathcal{S}^M$ with probability $P(C_\mathbf{n}|B_M) = P(C_\mathbf{n})/P(B_M)$ (where we note $C_\mathbf{n} \subset B_M$). This is the second factor on the right-hand side of Eq. (S8). The first factor, $P\left(A|C_\mathbf{n}\right)$, is the probability that $A$ occurs given that the population is in state $\mathbf{n}$.

In the stationary state, the distribution $P(C_{\mathbf{n}})$ is then given by the stationary distribution of the full model for a fixed value of $p_S$. Results for this distribution were obtained by Constable and Kokko in [1] for general values of $p_S$ ($p_S$ maps to the parameter $c$ in [1] via $c = 1 - p_S$). Using $P(C_{\mathbf{n}})$ we can express $P(B_M) = \sum_{n\in\mathcal{S}^M} P(C_{\mathbf{n}})$.

The rate $T^-_{M|p_S}$ is obtained from $P^-_{M|p_S}(\Delta t)$ as

$$T^-_{M|p_S} = \lim_{\Delta t\to 0} \frac{P^-_{M|p_S}(\Delta t)}{\Delta t}. \tag{S9}$$

The problem then reduces to calculating $P(A|C_{\mathbf{n}})$, for $\mathbf{n} \in \mathcal{S}^M$, i.e. the probability that a mating type becomes extinct in the next time interval $\Delta t$, if the composition of the population is currently $\mathbf{n}$.

As a first step, we find the overall rate with which $n_j \to n_j - 1$ for a fixed $j$ (and always assuming fixed $p_S$). This takes the form

$$\sum_{i\neq j} \mathcal{T}^{p_S}_{ij} = \frac{n_j}{2N}\left[(2-p_S)(N-n_j) - \frac{p_S}{N}\left(\sum_{k=1}^{M} n_k^2 - n_j^2\right)\right]. \tag{S10}$$

We write $\mathcal{F}^{p_S}_M(\mathbf{n})$ for overall rate of the event in which any entry $n_j = 1$ is reduced to zero. Summing over $j$ in Eq. (S10) and taking into account only contributions with $n_j = 1$, this is obtained as

$$\mathcal{F}^{p_S}_M(\mathbf{n}) = \frac{1}{2N}\left[(2-p_S)(N-1) + \frac{p_S}{N}\left(\sum_{k=1}^{M} n_k^2 - 1\right)\right]\left[\sum_{j=1}^{M}\delta_{n_j,1}\right]. \tag{S11}$$

In this expression $\delta_{n_j,1}$ denotes the Kronecker delta, i.e. $\delta_{n,1} = 1$ if $n = 1$ and $\delta_{n,1} = 0$ otherwise. The sum $\sum_{j=1}^{M}\delta_{n_j,1}$ is therefore the number of entries of $\mathbf{n}$ that are equal to one. With this, we obtain

$$P(A|C_{\mathbf{n}}) = \mathcal{F}^{p_S}_M(\mathbf{n})\Delta t. \tag{S12}$$

Putting everything together, we have

$$T^-_{M|p_S} = \frac{\displaystyle\sum_{\mathbf{n}\in\mathcal{S}^M} \mathcal{F}^{p_S}_M(\mathbf{n}) P^{\mathrm{st}}_{\mathbf{n}|p_S}}{\displaystyle\sum_{\mathbf{n}\in\mathcal{S}^M} P^{\mathrm{st}}_{\mathbf{n}|p_S}}. \tag{S13}$$

We stress again that this is for a fixed sex rate $p_S$ in the full model. The result is exact as long as we focus on long times (such that the population is in its stationary state). The denominator on the right-hand-side in Eq. (S13) is the stationary distribution of finding $M$ mating types for a given $p_S$, $P^{\mathrm{st}}_{M|p_S}$.

The sums in Eq. (S13) run over all $M-$compositions of $N$, $\mathbf{n} \in \mathcal{S}^M$. Using properties of sums over compositions it is possible to obtain a closed-form solution for $T^-_{M|p_S}$. We will describe this in detail below in Section S2.

##### B. Calculation of the effective rate $T^{+}_{M|p_S}$

To estimate $T^{+}_{M|p_S}$, we proceed in a similar way. We introduce

$$\begin{aligned} D = &\text{ the event in which the number of mating types increases from } M \\ &\text{ to } M+1 \text{ in the next } \Delta t, \end{aligned} \tag{S14}$$

and obtain

$$P^{+}_{M|p_S}(\Delta t) = \sum_{\mathbf{n}\in\mathcal{S}^M} P\left(D|C_{\mathbf{n}}\right) \frac{P(C_{\mathbf{n}})}{P(B_M)}. \tag{S15}$$

The probability $P(D|C_{\mathbf{n}})$ can be derived from the rate $\mathcal{T}_j^m$ in Eq. (S2). Assuming a fixed value of $p_S$, one gets

$$\begin{aligned} P(D|C_{\mathbf{n}}) &= \Delta t \sum_{j:n_j>1} \mathcal{T}_j^m \\ &= \Delta t \, \frac{m_g}{N} \left( N - \sum_{i=1}^{M} \delta_{n_i,1} \right). \end{aligned} \tag{S16}$$

The factor $N - \sum_{i=1}^{M} \delta_{n_i,1}$ on the right-hand side is the number of individuals in the population for which there is at least one other individual in the population with the same mating type (i.e, the sum excludes individuals who are 'singletons' in the population). Only mutation events of such individuals lead to an increase of the number of mating types. Introducing

$$\mathcal{G}_M(\mathbf{n}) = \frac{m_g}{N} \left( N - \sum_{i=1}^{M} \delta_{n_i,1} \right), \tag{S17}$$

we obtain

$$T^{+}_{M|p_S} = \frac{\sum_{\mathbf{n}\in\mathcal{S}^M} \mathcal{G}_M(\mathbf{n}) P^{\mathrm{st}}_{\mathbf{n}|p_S}}{\sum_{\mathbf{n}\in\mathcal{S}^M} P^{\mathrm{st}}_{\mathbf{n}|p_S}}. \tag{S18}$$

As for calculation of $T^{-}_{M|p_S}$, it is now necessary to carry out the sum over $M-$compositions of the integer $N$. We will describe in Section S2 how we do this.

#### S2. CLOSED-FORM SOLUTIONS OF $T^{-}_{M|p_S}$, $T^{+}_{M|p_S}$, AND $P^{\mathrm{st}}_{M|p_S}$

In this section, we present closed-form solutions of rates $T^{-}_{M|p_S}$, $T^{+}_{M|p_S}$, and stationary distribution $P^{\mathrm{st}}_{M|p_S}$ using the results derived in the previous section. The method we use is based on the number theory of partitions of integer numbers [2, 3].

As a first step we expand the terms inside the sums in Eqs. (S13) and (S18). We have

$$T^{-}_{M|p_S} = \frac{1}{2N}\left[(2-p_S)(N-1)+\frac{p_S}{N}\right]\frac{\sum_{\mathbf{n}\in\mathcal{S}^M}\left(\sum_{i=1}^{M}\delta_{n_i,1}\right)P^{\text{st}}_{\mathbf{n}|\sigma}}{\sum_{\mathbf{n}\in\mathcal{S}^M}P^{\text{st}}_{\mathbf{n}|\sigma}} - \frac{p_S}{2N^2}\frac{\sum_{\mathbf{n}\in\mathcal{S}^M}\left(\sum_{i=1}^{M}n_i^2\right)\left(\sum_{i=1}^{M}\delta_{n_i,1}\right)P^{\text{st}}_{\mathbf{n}|p_S}}{\sum_{\mathbf{n}\in\mathcal{S}^M}P^{\text{st}}_{\mathbf{n}|p_S}}, \tag{S19}$$

and

$$T^{+}_{M|p_S} = \frac{m_g}{N}\left[N - \frac{\sum_{\mathbf{n}\in\mathcal{S}^M}\left(\sum_{i=1}^{M}\delta_{n_i,1}\right)P^{\text{st}}_{\mathbf{n}|p_S}}{\sum_{\mathbf{n}\in\mathcal{S}^M}P^{\text{st}}_{\mathbf{n}|\sigma}}\right]. \tag{S20}$$

We notice that there are three sums involved:

$$\begin{aligned} S_1 &\equiv \sum_{\mathbf{n}\in\mathcal{S}^M}P^{\text{st}}_{\mathbf{n}|p_S}, \\ S_2 &\equiv \sum_{\mathbf{n}\in\mathcal{S}^M}\left(\sum_{i=1}^{M}\delta_{n_i,1}\right)P^{\text{st}}_{\mathbf{n}|p_S}, \\ S_3 &\equiv \sum_{\mathbf{n}\in\mathcal{S}^M}\left(\sum_{i=1}^{M}n_i^2\right)\left(\sum_{i=1}^{M}\delta_{n_i,1}\right)P^{\text{st}}_{\mathbf{n}|p_S}. \end{aligned} \tag{S21}$$

Each of these depends on the value of $p_S$. For $p_S = 0$, it is not necessary to calculate the last sum as the term including it in Eq. (S19) vanishes. In what follows, we will analyse the relevant sums for general values of $p_S$. We do this first for the asexual environment ($p_S = 0$), and then for general $0 < p_S \leq 1$.

##### A. Asexual environment ($p_S = 0$)

The stationary distribution of the full model for $p_S = 0$ ($\sigma = A$) was obtained in [1] as

$$P^{\text{st}}_{\mathbf{n}|A} = \alpha \times \prod_{k=1}^{M}\frac{1}{n_k}, \tag{S22}$$

where $\alpha$ represents a suitable normalisation constant. We write $P^{\text{st}}_{\mathbf{n}|A} = P^{\text{st}}_{\mathbf{n}|p_S=0}$. This constant plays no role in the evaluation of the expressions on the right-hand sides of Eqs. (S13) and (S18), as $P^{\text{st}}_{\mathbf{n}|A}$ appears in the numerator and in the denominator.

##### 1. *Calculation of* $S_1 = \sum_{\mathbf{n}\in\mathcal{S}^M} P^{st}_{\mathbf{n}|A}$

The sum $S_1 = \sum_{\mathbf{n}\in\mathcal{S}^M} P^{\rm st}_{\mathbf{n}|A}$ takes the following form as a sum over compositions

$$S_1 = \alpha \sum_{\substack{n_1+\dots+n_M=N\\ n_i\geq 1}} \prod_{k=1}^{M} \frac{1}{n_k}. \tag{S23}$$

This sum has the form $\sum_{n_1+\dots+n_M=N,\, n_i\geq 1} \prod_{k=1}^{M} f(n_k)$, with $f(n_k) = 1/n_k$. For sums of this form there exists a closed-form solution. It is given by (see e.g. [4])

$$\sum_{\substack{n_1+\dots+n_M=N\\ n_i\geq 1}} \prod_{k=1}^{M} f(n_k) = \frac{1}{N!}\frac{d^N}{dx^N}\bigg|_{x=0} \left(\sum_{i\geq 1} f(i)x^i\right)^M, \tag{S24}$$

i.e., the sum is equal to the coefficient multiplying $x^N$ in the power-series expansion of $\left(\sum_{i\geq 1} f(i)x^i\right)^M$.

To simplify matters we introduce the partition function $p(x) = \sum_{i\geq 1} f(i)x^i$, where the sum extends over all values of $i$ for which the function $f(i)$ is defined. In the present case, $f(i) = 1/i$, and $i$ can take arbitrarily large integer values. We find $p(x) = -\log(1-x)$.

Using Cauchy's integral formula [5] the coefficient multiplying $x^N$ in the expression in Eq. (S24) is given by

$$\frac{1}{N!}\frac{d^N}{dx^N}\bigg|_{x=0} p(x)^M = \frac{1}{2\pi i}\oint_\gamma \frac{p(z)^M}{z^{N+1}}\mathrm{d}z, \tag{S25}$$

with $\gamma$ an appropriate rectifiable curve around the origin. We find a solution to this integral by virtue of the integral representation of the Stirling numbers [6]. One obtains

$$\sum_{\substack{n_1+\dots+n_M=N\\ n_i\geq 1}} \prod_{k=1}^{M} \frac{1}{n_k} = \frac{M!}{N!}\begin{bmatrix} N\\ M\end{bmatrix}, \tag{S26}$$

where $\begin{bmatrix} N\\ M\end{bmatrix}$ is the unsigned Stirling number of the first kind [7].

##### 2. *Calculation of* $S_2 = \sum_{\mathbf{n}\in\mathcal{S}^M} \left(\sum_{i=1}^M \delta_{n_i,1}\right) P^{st}_{\mathbf{n}|A}$

The sum $\sum_{\mathbf{n}\in\mathcal{S}^M} \left(\sum_{i=1}^M \delta_{n_i,1}\right) P^{\rm st}_{\mathbf{n}|A}$ becomes

$$S_2 = \alpha \sum_{\substack{n_1+\dots+n_M=N\\ n_i\geq 1}} \left(\sum_{i=1}^{M} \delta_{n_i,1}\right) \prod_{k=1}^{M} \frac{1}{n_k}. \tag{S27}$$

Expanding the sum over the terms $\delta_{n_i,1}$, we find

$$\begin{aligned} S_2 &= \alpha\left(\sum_{\substack{n_1+\dots+n_M=N\\ n_i\geq 1}} \delta_{n_1,1}\prod_{k=1}^{M}\frac{1}{n_k} + \sum_{\substack{n_1+\dots+n_M=N\\ n_i\geq 1}} \delta_{n_2,1}\prod_{k=1}^{M}\frac{1}{n_k} + \dots + \sum_{\substack{n_1+\dots+n_M=N\\ n_i\geq 1}} \delta_{n_M,1}\prod_{k=1}^{M}\frac{1}{n_k}\right) \\ &= \alpha\left(\sum_{\substack{n_2+\dots+n_M=N-1\\ n_i\geq 1}} \prod_{k=2}^{M}\frac{1}{n_k} + \sum_{\substack{n_1+n_3+\dots+n_M=N-1\\ n_i\geq 1}} \frac{1}{n_1}\prod_{k=3}^{M}\frac{1}{n_k} + \dots + \sum_{\substack{n_1+\dots+n_{M-1}=N-1\\ n_i\geq 1}} \prod_{k=1}^{M-1}\frac{1}{n_k}\right) \\ &= \alpha M\left(\sum_{\substack{n_1+\dots+n_{M-1}=N-1\\ n_i\geq 1}} \prod_{k=1}^{M-1}\frac{1}{n_k}\right). \end{aligned} \tag{S28}$$

In the first line, we have applied the Kronecker delta functions $\delta_{n_i,1}$. In the second line, we have relabelled the $n_i$ so that each term takes the same form.

The resulting sum over compositions in the last line of Eq. (S28) is the same as the one in Eq. (S26), but with $N$ and $M$ replaced by $N-1$ and $M-1$, respectively. Therefore,

$$\sum_{\substack{n_1+\dots+n_M=N\\ n_i\geq 1}} \left(\sum_{i=1}^{M}\delta_{n_i,1}\right)\prod_{k=1}^{M}\frac{1}{n_k} = \frac{M!}{(N-1)!}\begin{bmatrix} N-1 \\ M-1 \end{bmatrix}. \tag{S29}$$

##### 3. Rates $T^-_{M|A}$ and $T^+_{M|A}$, and stationary distribution $P^{st}_{M|A}$

Substituting these results into Eqs. (S19) and (S20), we find

$$T^-_{M|A} = (N-1)\frac{\begin{bmatrix} N-1 \\ M-1 \end{bmatrix}}{\begin{bmatrix} N \\ M \end{bmatrix}}, \tag{S30}$$

and

$$T^+_{M|A} = m_g(N-1)\frac{\begin{bmatrix} N-1 \\ M \end{bmatrix}}{\begin{bmatrix} N \\ M \end{bmatrix}}. \tag{S31}$$

We stress again that $\begin{bmatrix} N-1 \\ M-1 \end{bmatrix}$ and $\begin{bmatrix} N \\ M \end{bmatrix}$ are not binomial coefficients, but instead unsigned Stirling numbers of the first kind.

Using the standard expression for the stationary distribution of two-species one-step birth-death processes (see Eq. (13) from the main text]), and after further algebra, we finally find

$$P^{st}_{M|A} = \frac{m_g^{M-1}}{(N-1)!}\frac{\begin{bmatrix} N \\ M \end{bmatrix}}{\binom{m_g+N-1}{m_g}}. \tag{S32}$$

##### B. Facultative sex ($0 < p_S < 1$) and sexual environment ($p_S = 1$)

We now consider values of $p_S > 0$. In this case the stationary distribution of the full model is given by (see [1] for details)

$$P^{\rm st}_{\mathbf{n}|p_S} = \alpha \prod_{k=1}^{M} \frac{1}{n_k\,(N_{p_S} - n_k)!}, \tag{S33}$$

with

$$N_{p_S} = N\left(\frac{2-p_S}{p_S}\right). \tag{S34}$$

In the case of purely sexual reproduction, $N_{p_S}$ reduces to $N_{p_S=1} = N$.

The constant $\alpha$ in Eq. (S33) again ensures normalisation, and is not important for the quantities we seek to calculate. The constant does not necessarily take the same value as the analogous constant in the previous section.

###### 1. Calculation of $S_1 = \sum_{\mathbf{n}\in\mathcal{S}^M} P^{st}_{\mathbf{n}|p_S}$

The sum $\sum_{\mathbf{n}\in\mathcal{S}^M} P^{\rm st}_{\mathbf{n}|p_S}$ can be written as sum over compositions

$$\sum_{\mathbf{n}\in\mathcal{S}^M} P^{\rm st}_{\mathbf{n}|p_S} = \alpha \sum_{\substack{n_1+\cdots+n_M=N\\ n_i\geq 1}} \prod_{k=1}^{M} \frac{1}{n_k(N_{p_S} - n_k)!}. \tag{S35}$$

As with the asexual environment, this sum has the form $\sum_{n_1+\cdots+n_M=N,\, n_i\geq 1}\prod_{k=1}^{M} f(n_k)$, where we now have $f(n_k) = 1/[n_k(N_{p_S}-n_k)!]$. Therefore,

$$\sum_{\mathbf{n}\in\mathcal{S}^M} P^{\rm st}_{\mathbf{n}|p_S} = \alpha\, \frac{1}{N!}\frac{d^N}{dx^N}\bigg|_{x=0} p(x)^M \tag{S36}$$

with $p(x) = \sum_{i\geq 1} f(i)x^i$. The index $i$ in the sum in $p(x)$ extends up to the maximum integer number for which $f(i)$ is defined. In the present case this maximum value is given by $\lfloor N_{p_S}\rfloor$ with $\lfloor\cdot\rfloor$ the floor function ($\lfloor x\rfloor$ is the largest integer lower than or equal to $x$). Thus $p(x)$ is a a finite sum. Defining $L = \lfloor N_{p_S}\rfloor$, we have

$$p(x)^M = \left(\sum_{i\geq 1}^{L} \frac{x^i}{i(N_{p_S}-i)!}\right)^M. \tag{S37}$$

Applying the multinomial theorem, i.e.,

$$(x_1 + x_2 + \cdots + x_m)^n = \sum_{\substack{k_1\geq 0,\ldots,k_m\geq 0\\ k_1+k_2+\cdots+k_m=n}} \binom{n}{k_1, k_2, \ldots, k_m} \prod_{i=1}^{m} x_i^{k_i}, \tag{S38}$$

the $M$-th power of the function $p(x)$ becomes

$$\begin{aligned} p(x)^M &= \sum_{\substack{k_1\geq 0,\dots,k_L\geq 0\\ k_1+k_2+\cdots+k_L=M}} \binom{M}{k_1,k_2,\dots,k_L} \prod_{i=1}^{L} \left(\frac{x}{i(N_{p_S}-i)!}\right)^{k_i} \\ &= \sum_{\substack{k_1\geq 0,\dots,k_L\geq 0\\ k_1+k_2+\cdots+k_L=M}} \binom{M}{k_1,k_2,\dots,k_L} \prod_{i=1}^{L} \left(\frac{1}{i(N_{p_S}-i)!}\right)^{k_i} x^{k_1+2k_2+\cdots+Lk_L}, \end{aligned} \tag{S39}$$

where the sum extends over all the partitions of $M$, i.e., to all the possible ways of expressing $M$ as a sum of non-positive integers (i.e., we allow them also to take zero values). This means we allow the number of parts, i.e., the positive summands in a partition, to vary in the sum evaluation.

Based on Eq. (S36), we now need to extract the coefficient multiplying $x_N$ in the expression in Eq. (S39). We therefore need to identify all terms $x^{k_1+2k_2+\cdots+Lk_L}$ in Eq. (S39) such that $k_i \geq 0$, $k_1+2k_2+\cdots+Lk_L = N$ and $k_1+\cdots+k_L = M$. The sequences $k_1,\dots,k_L$ that satisfy these two conditions represent compositions of $N$ into $M$ positive parts (i.e., with a fixed number $M$ of summands) [8]. For fixed $M$, there can be no parts greater than $N-M+1$, so necessarily $k_{N-M+2} = \cdots = k_L = 0$. Therefore,

$$\begin{aligned} \frac{1}{N!}\frac{d^N}{dx^N}\bigg|_{x=0} p(x)^M = &\sum_{\substack{k_1+k_2+\cdots+k_{N-M+1}=M\\ k_1+2k_2+\cdots+(N-M+1)k_{N-M+1}=N}} \binom{M}{k_1,k_2,\dots,k_{N-M+1}} \prod_{i=1}^{N-M+1} \left(\frac{1}{i(N_{p_S}-i)!}\right)^{k_i} \\ =&\frac{M!}{N!} \sum_{\substack{k_1+k_2+\cdots+k_{N-M+1}=M\\ k_1+2k_2+\cdots+(N-M+1)k_{N-M+1}=N}} \binom{N}{k_1,k_2,\dots,k_{N-M+1}} \prod_{i=1}^{N-M+1} \left(\frac{1}{i(N_{p_S}-i)!}\right)^{k_i} \\ =&\frac{M!}{N!} B_{N,M}(x_1,\dots,x_{N-M+1}), \end{aligned}$$

where $B_{k,\ell}(x_1,\dots,x_{k-\ell+1})$ is the incomplete Bell polynomial [8] evaluated at $x_i = (i-1)!/(N_{p_S}-i)!$ for $i \in \{1,2,\dots,k-\ell+1\}$. It is convenient to express $x_i$ in the form

$$x_i = (i-1)!(N_{p_S}-1)_{i-1} x_1, \tag{S40}$$

with $(N_{p_S}-1)_{i-1}$ the falling factorial of $(N_{p_S}-1)$ with respect to $(i-1)$. Next, we use the following relation [8]

$$B_{k,\ell}(ax_1,\dots,ax_{k-\ell+1}) = a^{\ell} B_{k,\ell}(x_1,\dots,x_{k-\ell+1}). \tag{S41}$$

We arrive at

$$\sum_{\substack{n_1+\cdots+n_M=N\\ n_i\geq 1}} \prod_{k=1}^{M} \frac{1}{n_k(N_{p_S}-n_k)!} = \frac{M!}{N!(N_{p_S}-1)!^M} B_{N,M}(y_1,\dots,y_{N-M+1}), \tag{S42}$$

with $y_i = (i-1)!(N_{p_S}-1)_{i-1}$.

##### 2. *Calculation of* $S_2 = \sum_{\mathbf{n}\in\mathcal{S}^M}\left(\sum_{i=1}^{M}\delta_{n_i,1}\right)P^{st}_{\mathbf{n}|p_S}$

To calculate $\sum_{\mathbf{n}\in\mathcal{S}^M}\left(\sum_{i=1}^{M}\delta_{n_i,1}\right)P^{\text{st}}_{\mathbf{n}|p_S}$, we proceed in the same way as we did for the asexual environment (see Section S2 A 2). This sum has the form

$$S_2 = \alpha \sum_{\substack{n_1+\cdots+n_M=N\\ n_i\geq 1}} \left(\sum_{i=1}^{M}\delta_{n_i,1}\right)\prod_{k=1}^{M}\frac{1}{n_k(N_{p_S}-n_k)!}. \tag{S43}$$

Expanding the sum over $i$ this yields

$$\begin{aligned}
&\sum_{\substack{n_1+\cdots+n_M=N\\ n_i\geq 1}} \delta_{n_1,1}\prod_{k=1}^{M}\frac{1}{n_k(N_{p_S}-n_k)!}+\cdots+\sum_{\substack{n_1+\cdots+n_M=N\\ n_i\geq 1}} \delta_{n_M,1}\prod_{k=1}^{M}\frac{1}{n_k(N_{p_S}-n_k)!}\\
&=\sum_{\substack{n_2+\cdots+n_M=N-1\\ n_i\geq 1}} \frac{1}{(N_{p_S}-1)!}\prod_{k=2}^{M}\frac{1}{n_k(N_{p_S}-n_k)!}+\cdots+\sum_{\substack{n_1+\cdots+n_{M-1}=N-1\\ n_i\geq 1}} \frac{1}{(N_{p_S}-1)!}\prod_{k=1}^{M-1}\frac{1}{n_k(N_{p_S}-n_k)!}\\
&=\frac{M}{(N_{p_S}-1)!}\sum_{\substack{n_1+\cdots+n_{M-1}=N-1\\ n_i\geq 1}}\prod_{k=1}^{M-1}\frac{1}{n_k(N_{p_S}-n_k)!}.
\end{aligned}\tag{S44}$$

As before, in the first equality we have applied the functions $\delta_{n_i,1}$, while in the second step, we have relabelled the terms $n_i$ so that each sum takes the same form. The resulting sum is nothing but Eq. (S42) using $N-1$ and $M-1$ instead of $N$ and $M$ (without affecting $N_{p_S}$), respectively. Therefore,

$$\sum_{\substack{n_1+\cdots+n_M=N\\ n_i\geq 1}} \left(\sum_{i=1}^{M}\delta_{n_i,1}\right)\prod_{k=1}^{M}\frac{1}{n_k(N_{p_S}-n_k)!} = \frac{M!}{(N-1)!(N_{p_S}-1)!^M}B_{N-1,M-1}(y_1,\ldots,y_{N-M+1}), \tag{S45}$$

with $y_i=(i-1)!(N_{p_S}-1)_{i-1}$.

##### 3. *Calculation of* $S_3 = \sum_{\mathbf{n}\in\mathcal{S}^M}\left(\sum_{i=1}^{M}n_i^2\right)\left(\sum_{i=1}^{M}\delta_{n_i,1}\right)P^{st}_{\mathbf{n}|p_S}$

This sum takes the form

$$S_3 = \sum_{\substack{n_1+\cdots+n_M=N\\ n_i\geq 1}} \left(\sum_{i=1}^{M}\delta_{n_i,1}\right)\frac{n_1^2+\cdots+n_M^2}{n_1(N_{p_S}-n_1)!\cdots n_M(N_{p_S}-n_M)!}. \tag{S46}$$

Expanding the sum over index $i$, and relabelling $n_i$ appropriately, yields

$$S_3 = \frac{M}{(N_{ps}-1)!}\left( \sum_{\substack{n_1+\cdots+n_{M-1}=N-1 \\ n_i \geq 1}} \prod_{k=1}^{M-1} \frac{1}{n_k(N_{ps}-n_k)!} + \sum_{\substack{n_1+\cdots+n_{M-1}=N-1 \\ n_i \geq 1}} \frac{n_1^2+\cdots+n_{M-1}^2}{\prod_{k=1}^{M-1} n_k(N_{ps}-n_k)!} \right) \tag{S47}$$

The first sum inside the brackets is of the same form as the expression in Eq. (S42), but with $N$ and $M$ replaced by $N-1$ and $M-1$, respectively, without affecting $N_{ps}$. Therefore we have

$$\sum_{\substack{n_1+\cdots+n_{M-1}=N-1 \\ n_i \geq 1}} \prod_{k=1}^{M-1} \frac{1}{n_k(N_{ps}-n_k)!} = \frac{(M-1)!}{(N-1)!(N_{ps}-1)!^{M-1}} B_{N-1,M-1}(y_1,\ldots,y_{N-M+1}). \tag{S48}$$

To calculate the second sum inside the brackets on the right-hand side of Eq. (S47), we note that by symmetry the $M-1$ terms resulting when expanding the numerator $n_1^2+\cdots+n_{M-1}^2$ are all equal after the sum over compositions is carried out. In other words

$$\sum_{\substack{n_1+\cdots+n_{M-1}=N-1 \\ n_i \geq 1}} \frac{n_1^2+\cdots+n_{M-1}^2}{\prod_{k=1}^{M-1} n_k(N_{ps}-n_k)!} = (M-1) \sum_{\substack{n_1+\cdots+n_{M-1}=N-1 \\ n_i \geq 1}} \frac{n_1}{(N_{ps}-n_1)!} \frac{1}{\prod_{k=2}^{M-1} n_k(N_{ps}-n_k)!}. \tag{S49}$$

This leads to

$$\sum_{\substack{n_1+\cdots+n_{M-1}=N-1 \\ n_i \geq 1}} \frac{n_1^2+\cdots+n_{M-1}^2}{\prod_{k=1}^{M-1} n_k(N_{ps}-n_k)!} = (M-1) \sum_{n_1=1}^{N-M+1} \frac{n_1}{(N_{ps}-n_1)!} \left( \sum_{\substack{n_2+\cdots+n_{M-1}=N-1-n_1 \\ n_i \geq 1}} \frac{1}{\prod_{k=2}^{M-1} n_k(N_{ps}-n_k)!} \right). \tag{S50}$$

The sum over $n_1$ ranges from the minimum to the maximum value $n_1$ can take in the compositions. The sum inside the brackets is of the same form as the expression in Eq. (S42) ,with $N$ and $M$ replaced by $N-1-n_1$ and $M-2$, respectively, without affecting $N_{ps}$. We therefore have

$$\sum_{\substack{n_1+\cdots+n_{M-1}=N-1 \\ n_i \geq 1}} \frac{n_1^2+\cdots+n_{M-1}^2}{\prod_{k=1}^{M-1} n_k(N_{ps}-n_k)!} = \sum_{n_1=1}^{N-M+1} \frac{n_1}{(N_{ps}-n_1)!} \frac{(M-1)!}{(N-1-n_1)!(N_{ps}-1)!^{M-2}} B_{N-1-n_1,M-2}(y_1,\ldots,y_{N-n_1-M+2}), \tag{S51}$$

with $y_i = (i-1)!(N_{ps}-1)_{i-1}$.

Putting everything together, we finally arrive at

$$\sum_{\substack{n_1+\cdots+n_M=N\\ n_i\geq 1}} \left(\sum_{i=1}^{M} \delta_{n_i,1}\right) \frac{n_1^2+\cdots+n_M^2}{n_1(N_{p_S}-n_1)!\cdots n_M(N_{p_S}-n_M)!} =$$

$$\frac{M!}{(N_{p_S}-1)!^{M-1}} \left( \frac{1}{(N-1)!(N_{p_S}-1)!} B_{N-1,M-1}(y_1,\ldots,y_{N-M+1}) + \sum_{n_1=1}^{N-M+1} \frac{n_1}{(N_{p_S}-n_1)!} \frac{1}{(N-1-n_1)!} B_{N-1-n_1,M-2}(y_1,\ldots,y_{N-n_1-M+2}) \right), \tag{S52}$$

with $y_i = (i-1)!(N_{p_S}-1)_{i-1}$.

###### 4. *Rates $T^-_{M|p_S}$ and $T^+_{M|p_S}$, and stationary distribution $P^{st}_{M|p_S}$*

Putting the different results together, we find for fixed rate of sexual reproduction $p_S > 0$:

$$T^-_{M|p_S} = \frac{1}{2}(2-p_S)(N-1)\frac{B_{N-1,M-1}(y_1,\ldots,y_{N-M+1})}{B_{N,M}(y_1,\ldots,y_{N-M+1})} - \frac{p_S}{2N}\frac{(N-1)!(N_{p_S}-1)!}{B_{N,M}(y_1,\ldots,y_{N-M+1})}\sum_{n_1=1}^{N-M+1}\frac{n_1}{(N_{p_S}-n_1)!}\frac{B_{N-1-n_1,M-2}(y_1,\ldots,y_{N-n_1-M+2})}{(N-1-n_1)!}, \tag{S53}$$

and

$$T^+_{M|p_S} = m_g\left(1-\frac{B_{N-1,M-1}(y_1,\ldots,y_{N-M+1})}{B_{N,M}(y_1,\ldots,y_{N-M+1})}\right). \tag{S54}$$

Recall we have $B_{k,\ell} = B_{k,\ell}(y_1,\ldots,y_{k-\ell+1})$ with $y_i = (i-1)!(N_{p_S}-1)_{i-1}$. From these rates we can construct the stationary distribution $P^{\mathrm{st}}_{M|p_S}$ using the standard formula for birth-death processes (see Eq. (13)).

We recall from Section III B in the main paper that the rates $T^-_{M|p_S}$ and $T^+_{M|p_S}$ from Eqs. (S53) and (S54) correspond to the rates in the fast switching limit, i.e., $T^{-,\mathrm{fast}}_M$ and $T^{-,\mathrm{slow}}_M$. As explained in there, in this limit the system behaves as if there was a fixed environment with sex rate $p_S$.

##### C. Alternative expressions of rates $T^-_{M|p_S}$ and $T^+_{M|p_S}$ for cases $0 < p_S \leq 1$

In Section S2 B we derived analytical closed-form expressions of rates $T^-_{M|p_S}$ and $T^+_{M|p_S}$ for $0 < p_S \leq 1$. The results are exact in the stationary state, but the evaluation of these expressions may be computationally costly for large population sizes $N$. We therefore proceed to derive equivalent expressions which are easier to evaluate numerically.

For that, we start from Eqs. (S19) and (S20) and express the sums over compositions as standard non-partition sums, i.e., as sums of indexed numbers with defined lower and upper bounds of summation. Denoting by $P$ the number of entries higher than 1 of a given state $\mathbf{n}$, we first express each sum over compositions in Eqs. (S19) and (S20) as sums over $P$. The sum $S_1$ (see Eq. (S21)) can be written in the form

$$\sum_{\mathbf{n}\in\mathcal{S}^M} P^{\text{st}}_{\mathbf{n}|p_S} = \sum_{P=0}^{P_{\max}} S^P_{M,p_S}(N-M), \tag{S55}$$

with a suitable $S^P_{M,p_S}(N-M)$ (see below), and where $P_{\max}$ is the maximum number of mating types with strictly more than one individual. For a given $N$ and $M$ this is

$$P_{\max} = \begin{cases} 0 & \text{if } N = M \\ N-M & \text{if } N/2 \leq M < N \\ M & \text{if } M < N/2. \end{cases} \tag{S56}$$

Notice that when $N = M$ then there will necessarily be exactly one individual of each mating type, so $P_{\max} = 0$.

The term $S^P_{M,p_S}(N-M)$ in Eq. (S55) represents the sum of $P^{\text{st}}_{\mathbf{n}|p_S}$ over compositions with $P$ elements higher than one for a given $N$ and $M$, i.e.,

$$S^P_{M,p_S} \equiv \sum_{\substack{n_1+\cdots+n_M=N \\ |G|=P \\ G=\{n_i>1:1\leq i\leq M\}}} P^{\text{st}}_{\mathbf{n}|p_S}, \tag{S57}$$

This can be expressed as a recursive sum of the form

$$S^P_{M,p_S}(Q) = (N_{p_S}-1)!\binom{M-P+1}{P}\sum_{k_P=2}^{Q-P+2}\frac{1}{k_P(N_{p_S}-k_P)!}S^{P-1}_{M,p_S}(Q+1-k_P), \tag{S58}$$

with base cases

$$S^1_{M,p_S}(Q) = \frac{M}{(N_{p_S}-1)!^{M-1}}\frac{1}{(Q+1)(N_{p_S}-Q-1)!} \quad \text{and} \quad S^0_{M,p_S}(Q) = \frac{1}{(N_{p_S}-1)!^M}\delta_{Q,0}. \tag{S59}$$

The procedure to derive the previous expressions is laborious but straightforward. The strategy for this is to sort the $M-$compositions $\mathbf{n}$ in $\sum_{\mathbf{n}\in\mathcal{S}^M} P^{\text{st}}_{\mathbf{n}|p_S}$ such that we sum them starting from those compositions which contain entries with the lowest values to those with the highest values. Then we identify which compositions contain the same elements and regroup them.

Following with the rest of the sums over compositions in Eqs. (S19) and (S20), one finds hat the sum $S_2$ takes the form

$$\sum_{\mathbf{n}\in\mathcal{S}^M}\left(\sum_{i=1}^{M}\delta_{n_i,1}\right)P^{\text{st}}_{\mathbf{n}|p_S} = \sum_{P=0}^{P_{\max}}(M-P)\times S^P_{M,p_S}(N-M), \tag{S60}$$

as $M-P$ is the number of entries equal to one for state $\mathbf{n}$. Following the same logic as with $S^P_{M,p_S}$, the sum $S_3$ becomes

$$\sum_{\mathbf{n}\in\mathcal{S}^M}\left(\sum_{i=1}^{M} n_i^2\right)\left(\sum_{i=1}^{M}\delta_{n_i,1}\right)P^{\rm st}_{\mathbf{n}|p_S}=\sum_{P=0}^{P_{\max}}(M-P)\times\Gamma^P_{M,p_S}(N-M), \tag{S61}$$

where the term $\Gamma^P_{M,p_S}(N-M)$ represents the sum of $\left(\sum_{i=1}^{M} n_i^2\right)P^{\rm st}_{\mathbf{n}|p_S}$ over compositions with $P$ elements higher than one. This is

$$\Gamma^P_{M,p_S}(Q)=\binom{M}{P}\frac{1}{(N_{p_S}-1)!^{M-P}}\sum_{k_P=2}^{Q-P+2}\frac{1}{k_P(N_{p_S}-k_P)!}\Lambda^{P-1}_{p_S}(M-1+k_P^2,N-M+1-k_P), \tag{S62}$$

with

$$\Lambda^P_{p_S}(L,Q)=\sum_{k_P=2}^{Q+2-P}\frac{1}{k_P(N_{p_S}-k_P)!}\Lambda^{P-1}_{p_S}(L-1+k_P^2,Q+1-k_P), \tag{S63}$$

with base cases

$$\Gamma^1_{M,p_S}(Q)=\frac{M}{(N_{p_S}-1)!^{M-1}}\Lambda^1_{p_S}(M,Q),\quad \Lambda^1_{p_S}(L,Q)=\frac{L-1+(Q+1)^2}{(Q+1)(N_{p_S}-Q-1)!}, \tag{S64}$$

and

$$\Gamma^0_{M,p_S}(Q)=\frac{M}{(N_{p_S}-1)!^{M}}\Lambda^0_{p_S}(M,Q),\quad \Lambda^0_{p_S}(L,Q)=\delta_{Q,0}. \tag{S65}$$

Putting everything together in Eqs. (S19) and (S20), we finally find

$$\begin{aligned}T^-_{M|p_S}=&\frac{1}{2N}\left[(2-p_S)(N-1)+\frac{p_S}{N}\right]\frac{\sum_{P=0}^{P_{\max}}(M-P)\times S^P_{M,p_S}(N-M)}{\sum_{P=0}^{P_{\max}}S^P_{M,p_S}(N-M)}\\&-\frac{p_S}{2N^2}\frac{\sum_{P=0}^{P_{\max}}(M-P)\times\Gamma^P_{M,p_S}(N-M)}{\sum_{P=0}^{P_{\max}}S^P_{M,p_S}(N-M)},\end{aligned} \tag{S66}$$

and

$$T^+_{M|p_S}=\frac{m_g}{N}\left(N-\frac{\sum_{P=0}^{P_{\max}}(M-P)\times S^P_{M,p_S}(N-M)}{\sum_{P=0}^{P_{\max}}S^P_{M,p_S}(N-M)}\right). \tag{S67}$$

##### D. Comparison against numerical simulations with fixed environment

In Figure S1 we compare the predictions for the rates $T^-_{M|p_S}$ and $T^+_{M|p_S}$ in Eqs. (S53) and (S54) against numerical simulations for different values of $p_S$ with $0 < p_S < 1$. Cases with $p_S = 0$ and $p_S = 1$ are displayed in Figure 3 in the main paper. A comparison of theory and numerical simulations for the stationary distribution at fixed $p_S$ is shown in Figure S2.

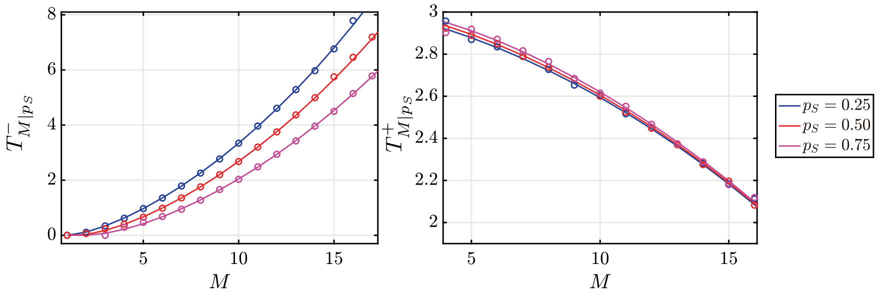


Figure S1. Rates $T^-_{M|p_S}$ and $T^+_{M|p_S}$ obtained from theory (solid lines) and numerical simulations of the full model (dotted lines) with fixed rate of sexual reproduction $p_S$. Parameters: $N = 30$, $m_g = 3.0$. Simulations were run up to time $t = 10^7$, with measurements starting $t = 10^6$ to ensure stationarity.

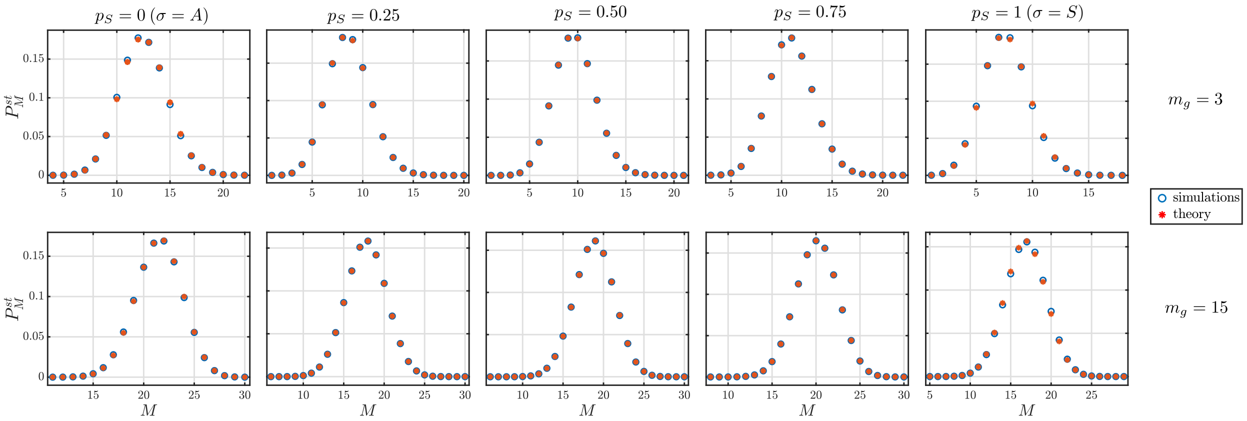


Figure S2. Stationary distribution $P^{\mathrm{st}}_{M|p_S}$ for fixed rate of sexual reproduction $p_S$ for population size $N = 30$ with $m_g = 3$ (upper row) and $m_g = 15$ (lower row). The theoretical predictions are given by the results presented in Sections S2 A 3 and S2 B 4. Numerical simulations are of the full model. Simulations were run up to time $t = 10^7$, with measurements starting $t = 10^6$ to ensure stationarity.

##### E. Comparison against numerical simulations with switching environments

The previous results assume a fixed environment with sex rate $p_S$. When introducing switching between environments $\sigma = S$ and $\sigma = A$, our predictions then may differ in certain circumstances. This discrepancy comes from the fact that rates $T^{\pm}_{M|\sigma}$ are calculated from the distribution of vector states $P^{\mathrm{st}}_{\mathbf{n}|\sigma}$ from the full model for a fixed environment (as shown in Section S1). In the case of switching environments the effective birth and death rates become $T^{\pm}_{M,\sigma}$, which specify the rate of transitions $(M, \sigma) \to (M \pm 1, \sigma)$. In this case $\sigma$ is not fixed as in rates $T^{\pm}_{M|\sigma}$. Following the same idea presented in Section S1, rates $T^{\pm}_{M,\sigma}$ would need to be obtained from $P^{\rm st}_{\mathbf{n},\sigma}$, i.e., the stationary distribution of states $(n, \sigma)$ in the full model with switching environments. Obtaining $P^{\rm st}_{\mathbf{n},\sigma}$, however, is beyond the scope of the present work.

In this section we explore this difference when varying the fraction of time spent in the sexual environment, $p_S$. As explained in Section III C, the fact of using rates $T^{\pm}_{M|\sigma}$ in the generator-matrix approach has consequences in the prediction of the stationary distribution $P^{\rm st}_M$ when using this approach for intermediate and fast-switching regimes. We provide more details of this in the next section.

In Figure S3, we illustrate a comparison of $T^{\pm}_{M,S}$ (obtained from simulations) against the theoretical prediction of $T^{\pm}_{M|S}$. As shown, the prediction gets better as the fraction time spent in environment $\sigma = S$ is higher (i.e., as $p_S$ increases). Conversely, we compare $T^{\pm}_{M,A}$ and $T^{\pm}_{M|A}$ in Figure S4, showing that the prediction gets worse when increasing $p_S$.

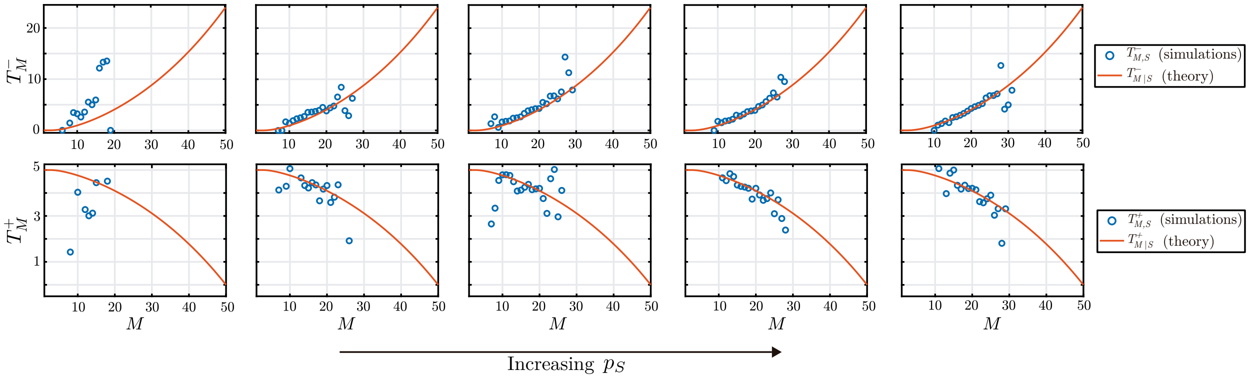


Figure S3. Comparison between $T^{\pm}_{M,S}$ (simulations) and $T^{\pm}_{M|S}$ (theory). We fix the average time cycle $\tau = 2$ generations throughout and vary the fraction of time spent in the sexual environment, $p_S$. From right to left panels, we set for each column $p_S = 0.01, 0.25, 0.50, 0.75, 0.99$. Remaining parameters: $N = 50$, $m_g = 5$.

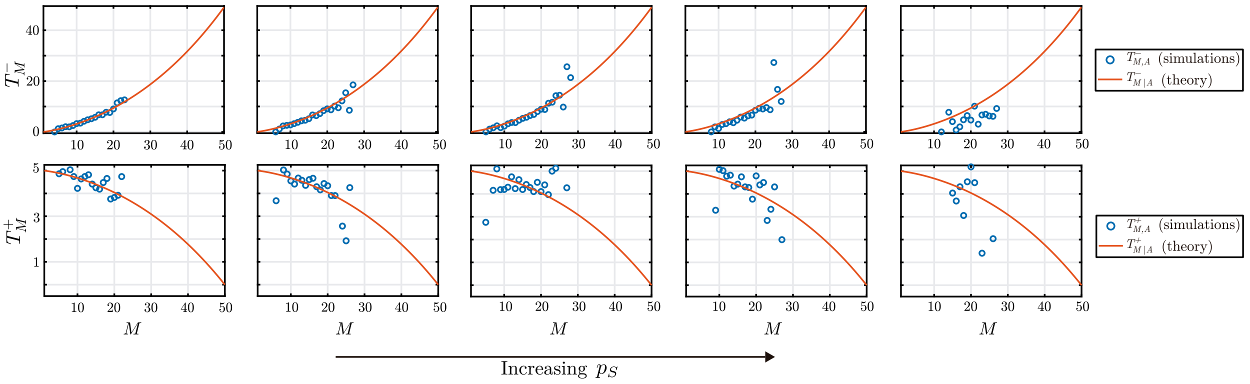


Figure S4. Comparison between $T^{\pm}_{M,A}$ (simulations) and $T^{\pm}_{M|A}$ (theory). We fix the average time cycle $\tau = 2$ generations throughout and vary the fraction of time spent in the sexual environment, $p_S$. From right to left panels, we set for each column $p_S = 0.01, 0.25, 0.50, 0.75, 0.99$. Remaining parameters: $N = 50$, $m_g = 5$.

### S3. MARKOV CHAIN GENERATOR MATRIX APPROACH

As explained in Section III C, the rates $T^{+}_{M,\sigma}$ and $T^{-}_{M,\sigma}$ used to construct the generator matrix $\underline{\underline{Q}}$ are those derived for situations in which the environment does not switch between states. In other words, we use the rates $T^{+}_{M|\sigma}$ and $T^{-}_{M|\sigma}$ ($\sigma = A, S$) obtained in Sections S1 A and S1 B of this Supplement ($\sigma = A$ corresponds to setting $p_S = 0$, and $\sigma = S$ to $p_S = 1$). Assuming a fixed value of $p_S$ is an approximation for the model in which the environment switches between the states with purely sexual and purely asexual reproduction. In this section, we explore the implications of this approximation.

#### A. Comparison against simulations

Figure S5 shows how the stationary distribution $P^{\rm st}_M$ obtained from the generator-matrix approach compares against numerical simulations. For the graphs shown, we have fixed the average period of one environmental switching cycle, $\tau$, while varying the average fraction spent in each environment. We show how the distribution compares to simulations for different values of $m_g$ for which, based on Figure 9, we expect to see a good or bad agreement. As shown, as $m_g$ increases the prediction from the generator-matrix approach becomes better. In Section III C 2 we discuss the reasons of this.

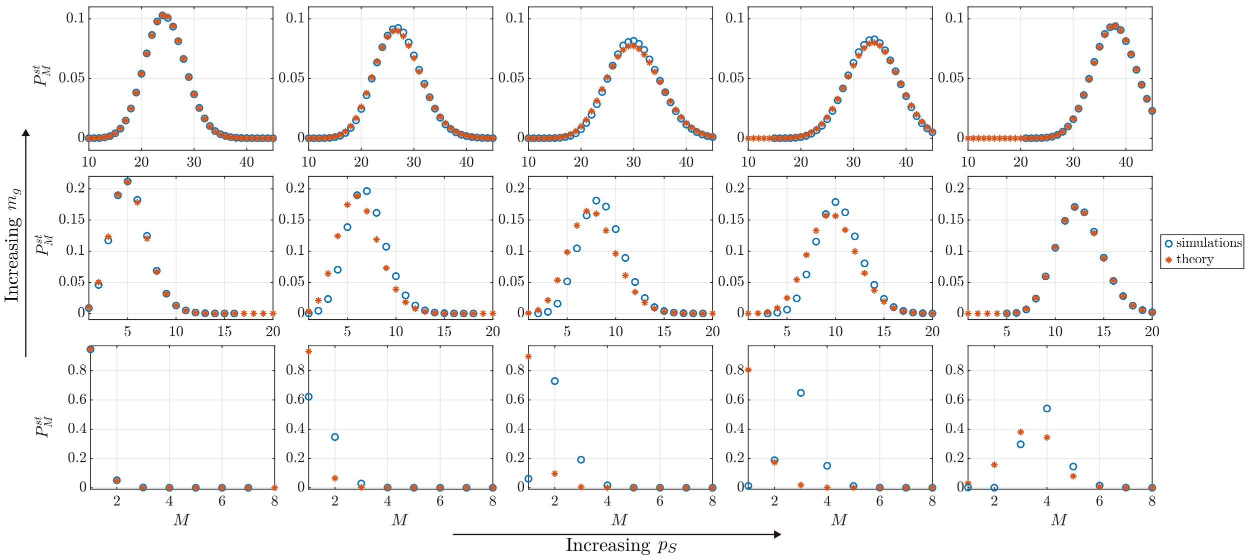


Figure S5. Stationary distribution $P^{\rm st}_M$ for $N = 100$ from numerical simulations of the full model with switching environment (open circles) and from the generator-matrix approach (asterisks). We fix the average cycle time to $\tau = 2$ generations throughout, and vary the fraction of time spent in the sexual environment, $p_S$, and the mutation rate, $m_g$. The values used (from left to right columns) are $p_S = 0.01, 0.25, 0.50, 0.75, 0.99$, and (from lower to upper rows) $m_g = 0.01, 1, 10$. Numerical simulations were conducted by time-averaging one single run up to time $t = 10^5$ generations, where the first $10^2$ units of time are ignored to ensure stationarity

##### B. Theoretical prediction of $P^{\mathrm{st}}_{M,\sigma}$ for each environment

Using the generator-matrix approach we can derive the stationary distribution $P^{\mathrm{st}}_{M,\sigma}$. We point out that this is not the same as $P^{\mathrm{st}}_{M|\sigma}$, instead $P^{\mathrm{st}}_{M,\sigma}$ is the joint distribution of finding the environment in state $\sigma$ and $M$ mating types in the population in a model with switching environment. The distribution $P^{\mathrm{st}}_{M,\sigma}$ cannot be calculated using the approaches for the slow switching and fast switching limits presented in Sections III A and III B.

As explained in Section III C 2, the first $N$ elements of $\underline{P}^{\mathrm{st}}$ are the $P^{\mathrm{st}}_{M,S}$, for $M = 1, \ldots, N$, while the second $N$ entries are $P^{\mathrm{st}}_{M,A}$ (see Eq. (23)). In Figure S6, we illustrate how the theoretical predictions for both distributions obtained from the generator-matrix approach behave for different values of $p_S$, i.e, when varying the fraction of time spent in each environment. These quantities cannot be obtained using the fast and slow switching approximations presented in Sections III A and III B. As shown, the amplitude of $P^{\mathrm{st}}_{M,S}$ increases with $p_S$, while the opposite occurs to $P^{\mathrm{st}}_{M,A}$. When $p_S = 0$ or $p_S = 1$ we see the distribution of only one environment, while when $p_S = 0.5$ the amplitude is similar in both distributions.

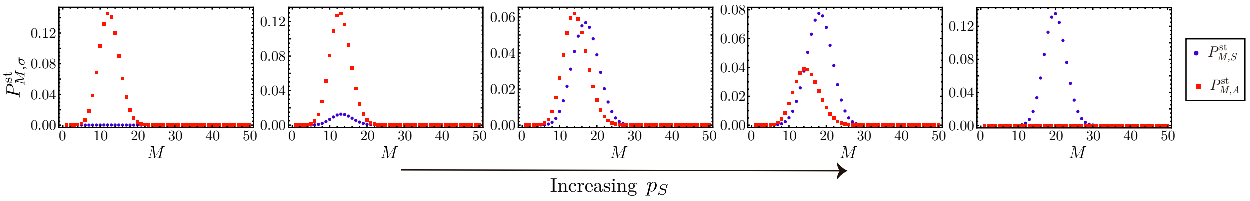


Figure S6. Predictions for the distributions $P^{\mathrm{st}}_{M,S}$ and $P^{\mathrm{st}}_{M,A}$ from the generator-matrix approach. Parameters $N = 50, m_g = 5$, and $\lambda_{A\to S} = 0.5$. The value for $p_S$ in the different panels is $0, 0.1, 0.5, 0.66, 1$ from left to right.

##### C. Transition between slow and fast switching regimes for low and high $p_S$

In Figure 7 of the main text, we showed how $P^{\mathrm{st}}_{M}$ behaves when varying the average time cycle $\tau$ for $p_S = 0.5$, i.e. when the environment spends equal fractions of time in each of the two states $A$ and $S$. The stationary distribution for $M$ the differs in the fast and slow switching regimes (for example the distribution can be bimodal for slow switching, but unimodal for fast switching). In Figure S7 we show how $P^{\mathrm{st}}_{M}$ behaves when the environment spends most of the time in either one of the two states. As shown, there is then no noticeable difference between the distributions in the slow and fast regimes. This is because one of the environments dominates, irrespective of the speed of environmental switching.

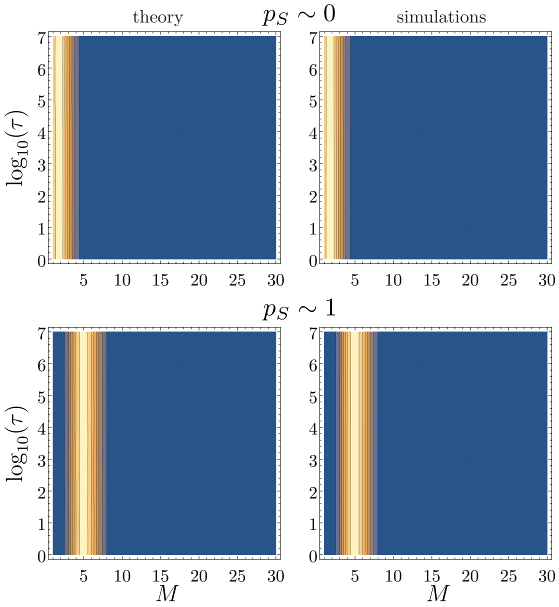


Figure S7. Stationary distribution $P_M^{\mathrm{st}}$ as a function of the cycle time $\tau$ obtained from the generator-matrix theoretical approach and simulations of the full model. The upper and lower rows represent cases of high and low $p_S$ with values $p_S = 0.999$ and $p_S = 0.001$, respectively. Parameters: $N = 30, m_g = 0.3$ and $\lambda_{A\to S} = 1$. Numerical simulations were conducted by time-averaging one single run up to time $t = 10^7$ generations, where the first $10^6$ units of time are ignored to ensure stationarity.

##### D. Analytical results from the generator-matrix approach

The theoretical approach in Section III C 2 of the main paper makes use of the generator matrix $\underline{\underline{Q}}$ to calculate the stationary distribution $\underline{P}^{\mathrm{st}}$. The evaluation of this is mostly numerical. We now show how analytical results can be derived from the generator-matrix approach in the slow and fast switching regimes. As we will see, this reproduces the results for the slow-switching limit in Section III A of the main paper, while the prediction from the generator matrix in the fast-switching limit is different from the result presented in Section III B for the full model. This disagreement originates from the use of the rates $T^-_{M|\sigma}$ and $T^+_{M|\sigma}$ in the generator-matrix approach. These rates are derived for fixed environments, and assuming a stationary state of the population in those environments, see Sections S1 and S1 B of this Supplement. These assumptions are not valid when the switching of the environment is fast, as the population can then not reach a stationary state between switches of the environmental state. However, as explained in Section III C 3, there are situations in which the fast-switching limit derived from the the generator-matrix approach approximates the predictions of the full model well (i.e., the limit presented in Section III B).

The analysis starts from the equation defining the stationary state of the reduced model,

$$\underline{P}^{\mathrm{st}}\underline{\underline{Q}} = 0. \tag{S68}$$

This is a linear $2N \times 2N$ system, and can be written out explicitly. For the components describing environmental state $\sigma = A$ we have

$$0 = T^-_{M-1|A}P^{\mathrm{st}}_{M-1,A} + T^-_{M+1|A}P^{\mathrm{st}}_{M+1,A} - (T^-_{M|A} + T^+_{M|A})P^{\mathrm{st}}_{M,A} + \lambda_{S\to A}P^{\mathrm{st}}_{M,S} - \lambda_{A\to S}P^{\mathrm{st}}_{M,A}, \tag{S69}$$

while for $\sigma = S$

$$0 = T^-_{M-1|S} P^{\rm st}_{M-1,S} + T^-_{M+1|S} P^{\rm st}_{M+1,S} - (T^-_{M|S} + T^+_{M|S}) P^{\rm st}_{M,S} + \lambda_{A\to S} P^{\rm st}_{M,A} - \lambda_{S\to A} P^{\rm st}_{M,S}. \quad \text{(S70)}$$

In these equations, $M$ takes values $1, \ldots, N$. At the boundaries we have $T^-_{1|\sigma} = 0$ and $T^+_{N|\sigma} = 0$ both for $\sigma = A$ and $\sigma = S$.

Subtracting Eq. (S70) from Eq. (S69) we find

$$T^-_{M|A} P^{\rm st}_{M,A} + T^-_{M|S} P^{\rm st}_{M,S} = T^+_{M-1|A} P^{\rm st}_{M-1,A} + T^+_{M-1|S} P^{\rm st}_{M-1,S}. \quad \text{(S71)}$$

Within the reduced model this relation holds for any combination of the environmental switching rates $\lambda_{A\to S}$ and $\lambda_{S\to A}$.

Before analysing the slow-switching and fast-switching regimes, we make further preparations. First, we recall that

$$\sum_{M=1}^{N} P^{\rm st}_{M,A} = 1 - p_S, \quad \text{and} \quad \sum_{M=1}^{N} P^{\rm st}_{M,S} = p_S, \quad \text{(S72)}$$

see Section II B of the main paper.

Secondly, by summing the first $M$ instances of Eqs. (S69) and (S69) we obtain

$$P^{\rm st}_{M,A} = \frac{T^+_{M-1|A}}{T^-_{M|A}} P^{\rm st}_{M-1,A} + \frac{\lambda_{S\to A}}{T^-_{M|A}} \sum_{M'=1}^{M-1} P^{\rm st}_{M',A} - \frac{\lambda_{A\to S}}{T^-_{M|A}} \sum_{M'=1}^{M-1} P^{\rm st}_{M',S} \quad \text{(S73)}$$

and

$$P^{\rm st}_{M,S} = \frac{T^+_{M-1|S}}{T^-_{M|S}} P^{\rm st}_{M-1,S} + \frac{\lambda_{A\to S}}{T^-_{M|S}} \sum_{M'=1}^{M-1} P^{\rm st}_{M',S} - \frac{\lambda_{S\to A}}{T^-_{M|S}} \sum_{M'=1}^{M-1} P^{\rm st}_{M',A}, \quad \text{(S74)}$$

respectively. In these relations we have expressed $P^{\rm st}_{M,\sigma}$ in terms of $P^{\rm st}_{M',\sigma}$ up to $M' = M - 1$.

##### 1. *Limit of slow environmental switching*

In the limit of slow switching, i.e., when $\lambda_{A\to S}, \lambda_{S\to A} \ll 1$, the dominant terms in Eqs. (S73) and (S74) are those not proportional to any of the switching rates, so that

$$P^{\rm st}_{M,\sigma} \approx \frac{T^+_{M-1|\sigma}}{T^-_{M|\sigma}} P^{\rm st}_{M-1,\sigma} \quad \text{(S75)}$$

both for $\sigma = A$ and $\sigma = S$. The rates $T^{\pm}_{M|\sigma}$ satisfy the following relation (see Eq. (13) in the main paper)

$$P^{\rm st}_{M|\sigma} = \frac{T^+_{M-1|\sigma}}{T^-_{M|\sigma}} P^{\rm st}_{M-1|\sigma}. \quad \text{(S76)}$$

From this we conclude

$$P^{\rm st}_{M,\sigma} \approx \frac{P^{\rm st}_{M|\sigma}}{P^{\rm st}_{M-1|\sigma}} P^{\rm st}_{M-1,\sigma} \tag{S77}$$

By applying this relation successively, one then sees that

$$P^{\rm st}_{M,\sigma} \approx \frac{P^{\rm st}_{M|\sigma}}{P^{\rm st}_{1|\sigma}} P^{\rm st}_{1,\sigma}. \tag{S78}$$

Summing on both sides over $M$ from 1 to $N$, and using $\sum_{M=1}^{N} P_{M|\sigma} = 1$, one finds

$$\sum_{M=1}^{N} P^{\rm st}_{M,\sigma} \approx \frac{P^{\rm st}_{1,\sigma}}{P^{\rm st}_{1|\sigma}}. \tag{S79}$$

This holds for $\sigma = A$ and $\sigma = S$. Using Eqs. (S72), these relations then become

$$(1-p_S) \approx \frac{P^{\rm st}_{1,A}}{P^{\rm st}_{1|A}}, \quad \text{and} \quad p_S \approx \frac{P^{\rm st}_{1,S}}{P^{\rm st}_{1|S}}. \tag{S80}$$

for $\sigma = A$ and $\sigma = S$, respectively. Finally, using this in Eq. (S78), we find

$$P^{\rm st}_{M,A} \approx (1-p_S) P^{\rm st}_{M|A}, \quad \text{and} \quad P^{\rm st}_{M,S} \approx p_S P^{\rm st}_{M|S}, \tag{S81}$$

which leads to

$$P^{\rm st}_{M} \approx (1-p_S) P^{\rm st}_{M|A} + p_S P^{\rm st}_{M|S}, \tag{S82}$$

which is the result in Section III A of the main paper.

##### *2. Limit of fast environmental switching*

In the limit of fast switching, the switching rates $\lambda_{A\to S}$ and $\lambda_{S\to A}$ take very large values ($\lambda_{A\to S}, \lambda_{S\to A} \gg 1$). The terms proportional to these rates in Eqs. (S69) and (S70) then dominate all other contributions in these equations. This leads to

$$\lambda_{A\to S} P^{\rm st}_{M,A} \approx \lambda_{S\to A} P^{\rm st}_{M,S}. \tag{S83}$$

Using $p_S = \lambda_{A\to S}/(\lambda_{A\to S} + \lambda_{S\to A})$ this can be written as

$$P^{\rm st}_{M,A} \approx \frac{(1-p_S)}{p_S} P^{\rm st}_{M,S}. \tag{S84}$$

This implies that the distributions $P_{M,A}$ and $P_{M,S}$ share the same mode for $M$ in the limit of fast environments.

Using Eq. (S84) in Eq. (S71) yields

$$\left(T^{-}_{M|A}\frac{(1-p_S)}{p_S}+T^{-}_{M|S}\right)P^{\rm st}_{M,S}\approx\left(T^{+}_{M-1|A}\frac{(1-p_S)}{p_S}+T^{+}_{M-1|S}\right)P^{\rm st}_{M-1,S}, \tag{S85}$$

and therefore,

$$\begin{aligned}P^{\rm st}_{M,S}&\approx\left(\frac{(1-p_S)T^{+}_{M-1|A}+p_ST^{+}_{M-1|S}}{(1-p_S)T^{-}_{M|A}+p_ST^{-}_{M|S}}\right)P^{\rm st}_{M-1,S},\\&=\frac{T^{+}_{M-1,\rm eff}}{T^{-}_{M,\rm eff}}P^{\rm st}_{M-1,S}.\end{aligned} \tag{S86}$$

This resembles the stationary distribution solution for fixed environments (see Eq. (13) in the main paper) with rates

$$T^{\pm}_{M,\rm eff}=(1-p_S)T^{\pm}_{M|A}+p_ST^{\pm}_{M|S}. \tag{S87}$$

This means that in the limit of fast environments the population behaves as if it were in an effective fixed environment with rates $T^{\pm}_{M,\rm eff}$.

By applying Eq. (S86) successively one sees that

$$P^{\rm st}_{M,S}\approx\frac{T^{+}_{M-1,\rm eff}\cdots T^{+}_{1,\rm eff}}{T^{-}_{M,\rm eff}\cdots T_{2,\rm eff}}P^{\rm st}_{1,S}, \tag{S88}$$

and, by using Eq. (S84), that

$$P^{\rm st}_{M,A}\approx\frac{(1-p_S)}{p_S}\frac{T^{+}_{M-1,\rm eff}\cdots T^{+}_{1,\rm eff}}{T^{-}_{M,\rm eff}\cdots T_{2,\rm eff}}P^{\rm st}_{1,S}. \tag{S89}$$

Finally, we obtain

$$P^{\rm st}_{M}\approx\frac{1}{p_S}\frac{T^{+}_{M-1,\rm eff}\cdots T^{+}_{1,\rm eff}}{T^{-}_{M,\rm eff}\cdots T_{2,\rm eff}}P^{\rm st}_{1,S}. \tag{S90}$$

The term $P^{\rm st}_{1,S}$ is a normalisation factor which is determined from normalisation ($\sum_{M=1}^{N}P^{\rm st}_{M}=1$). We find

$$P^{\rm st}_{1,S}=\left[\frac{1}{p_S}\left(1+\sum_{M=2}^{N}\prod_{i=2}^{M}\frac{T^{+}_{i-1,\rm eff}}{T^{-}_{i,\rm eff}}\right)\right]^{-1}. \tag{S91}$$

We remark that this prediction for the limit of fast environments is different from the result in Section III B in the main paper. In the main paper, the transition rates in the full model were approximated by the weighted average of the rates in each environment, with the weights given by the fraction spent in each environment, i.e.,

$$\mathcal{T}^{\rm fast}_{ij}=(1-p_S)\mathcal{T}^{A}_{ij}+p_S\mathcal{T}^{S}_{ij}. \tag{S92}$$

This approximation translates into

$$T_M^{\pm,\text{fast}} = (1-p_S)T^{\pm}_{M,A} + p_S T^{\pm}_{M,S} \tag{S93}$$

in the reduced model. Since we do not know how to calculate $T^{\pm}_{M,S}$ and $T^{\pm}_{M,A}$, we use $T^{\pm}_{M|S}$ and $T^{\pm}_{M|A}$ instead as input in the generator matrix $\underline{\underline{Q}}$. This leads to what we obtained above in Eq. (S87). Figure 9 in the manuscript explores the validity of this approximation. Nevertheless, we remark that although we do not calculate $T^{\pm}_{M,S}$ and $T^{\pm}_{M,A}$, we still are able to compute $T_M^{\pm,\text{fast}}$ (as explained in Section III B) by using the results from Section S2 B 4.

#### S4. SELECTIVE SWEEPS

In this section, we explore a model with additional selective sweeps, as described in Section IV of the main text. We present the theoretical formalism to derive the stationary distribution $P_M^{\text{st}}$ for this model, and compare the resulting predictions against numerical simulations. We consider situations with switching environments (so that reproduction switches between sexual and asexual), and the case of a fixed environment in which reproduction is only asexual. In the case of switching environments, selective sweeps can only occur in the asexual environment.

##### A. Switching environments

To estimate the distribution $P_M^{\text{st}}$ in the presence of selective sweeps we proceed using the method explained in Section III C 2 of the main text. We construct the generator matrix $\underline{\underline{Q}}$ and calculate the corresponding stationary state. Throughout this section the rates $T^{\pm}_{M|\sigma}$ are always those of the model without selective sweeps. Sweeps are accounted for separately in the analysis.

As in the main text, we write the matrix $\underline{\underline{Q}}$ in block form

$$\underline{\underline{Q}} = \begin{pmatrix} Q^{(A,A)} & Q^{(A,S)} \\ Q^{(S,A)} & Q^{(S,S)} \end{pmatrix}. \tag{S94}$$

Since selective sweeps occur only in the asexual environment, the block $Q^{(A,A)}$ is the only one affected by selective sweeps. Taking into account the process $M \xrightarrow{\nu} 1$, the block $Q^{(A,A)}$ becomes

$$Q^{(A,A)} = \begin{pmatrix} -(T^+_{1|A}+\lambda_{A\to S}) & T^+_{1|A} & 0 & \dots & \dots & 0 \\ T^-_{2|A}+\nu & -(T^+_{2|A}+T^-_{2|A}+\lambda_{A\to S}+\nu) & T^+_{2|A} & 0 & \dots & \vdots \\ \nu & T^-_{3|A} & \ddots & \ddots & \ddots & \vdots \\ \vdots & 0 & \ddots & \ddots & \ddots & 0 \\ \vdots & \vdots & \ddots & \ddots & \ddots & T^+_{N-1|A} \\ \nu & \dots & \dots & 0 & T^-_{N|A} & -(T^-_{N|A}+\lambda_{A\to S}+\nu) \end{pmatrix},$$

i.e., compared to the model without selective sweeps, we have added $\nu$ to the entries $Q^{(A,A)}_{M,1}$ and

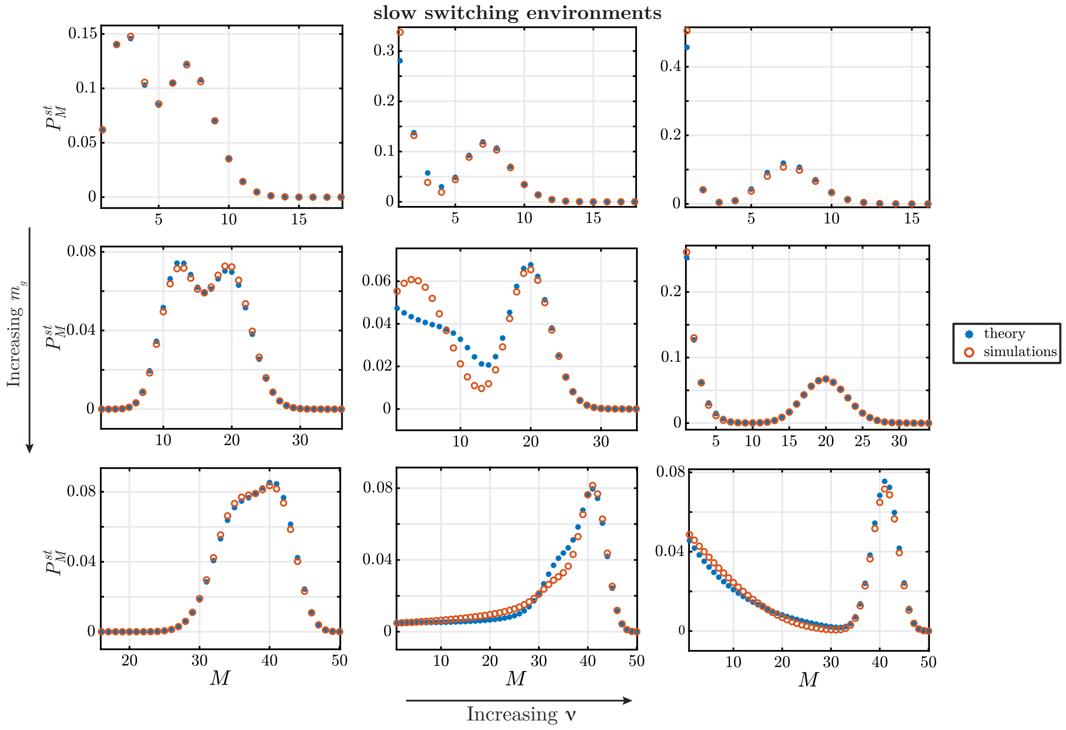


Figure S8. Stationary distribution of number of mating types in the model with selective sweeps and fast-switching environments. Theoretical prediction of $P_M^{\rm st}$ with selective sweeps as function of $M$ for slow switching environments, population size $N = 50$, for different values of $m_g$ and $\nu$. From upper to lower rows: $m_g = 0.5, 5, 50$. From left to right columns: $\nu = 0, 0.5, 5$. Switching rates used: $\lambda_{S\to A} = \lambda_{A\to S} = 5 \times 10^{-4}$.

$Q^{(A,A)}_{M,M}$ for $2 < M \le N$.

The stationary distribution $P_M^{\rm st}$ resulting from this generator matrix is an approximative result as the rates $T^{\pm}_{M|\sigma}$ are for fixed environments and do not account for selective sweeps. To take into account the effect of the selective sweeps on $T^{\pm}_{M|\sigma}$ it would be necessary to calculate first the stationary distribution $P^{\rm st}_{\mathbf{n},\sigma}$ from the full model, including selective sweeps. With this distribution one could then construct $T^{\pm}_{M|\sigma}$ in a similar way as explained in Section S1. It is difficult however to obtain $P^{\rm st}_{\mathbf{n},\sigma}$ for the full model with selective sweeps.

The generator-matrix approach still allows to capture important features that emerge from the inclusion of selective sweeps. In Figures S8 and S9, we illustrate the behaviour of $P_M^{\rm st}$ for slow switching and fast switching environments, respectively, as the parameters $m_g$ and $\nu$ are varied. Each of the graphs shows a horizontal cut in Figure 12 in the main text. We compare this against numerical simulations of the full model. In the simulations, the effect of selective sweeps is implemented by adding an extra transition $M \xrightarrow{\nu} 1$ in the asexual environment, independent of the state of the population. When such an event occurs, the population jumps to the state $M = 1$, and $n_1 = N$.

Figures S8 and S9 both show that, in general, the theoretical prediction is in good agreement with numerical simulations. As shown, the effect of selective sweeps is to increase the chance to find the system at $M = 1$, while mutations, on the contrary, have the effect of driving the population to higher numbers of mating types, reducing the probability of finding $M = 1$. When one of these effects dominates, our prediction shows good agreement with simulations. In such cases, the

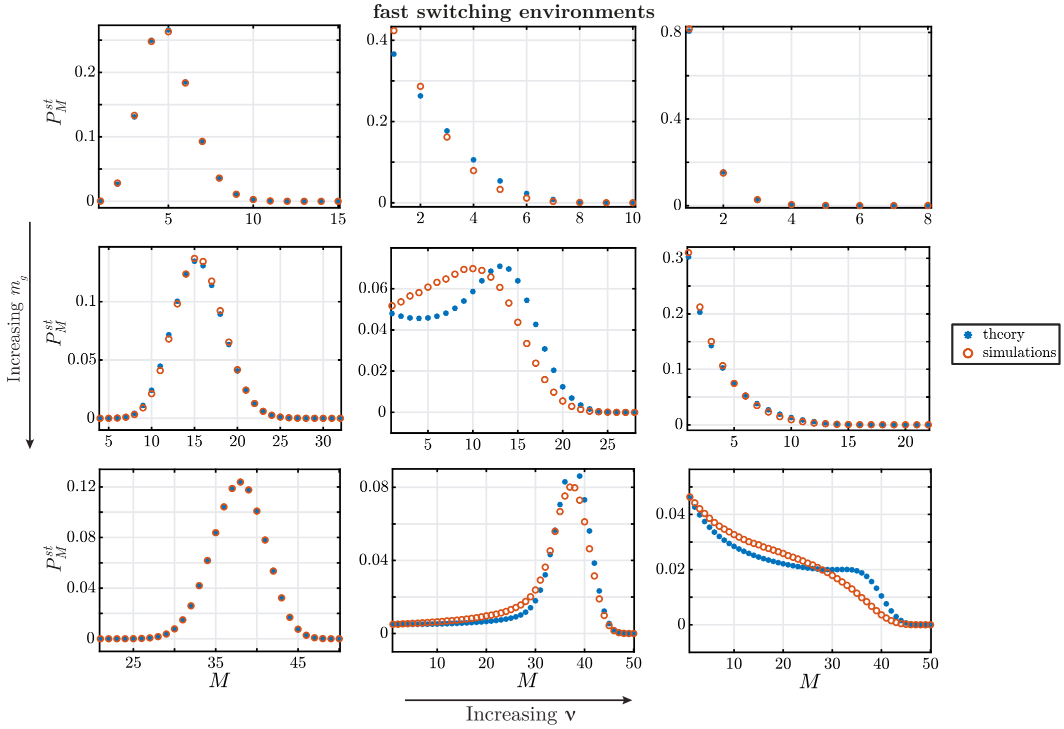


Figure S9. Stationary distribution of number of mating types in the model with selective sweeps and slowly switching environments. Theoretical prediction of $P_M^{\rm st}$ with selective sweeps as function of $M$ for fast switching environments, population size $N = 50$, for different values of $m_g$ and $\nu$. From upper to lower rows: $m_g = 0.5, 5, 50$. From left to right columns: $\nu = 0, 0.5, 5$. Switching rates used: $\lambda_{S\to A} = \lambda_{A\to S} = 5$.

distribution $P_M^{\rm st}$ exhibits a well-defined peak given by the dominant effect (at $M = 1$ when $\nu \gg m_g$, and at $M \gg 1$ when $m_g \gg \nu$). In situations like this, the assumption of ignoring selective sweeps on the calculation of rates $T^{\pm}_{M|\sigma}$ may become less relevant: if $\nu \gg m_g$, selective sweeps are dominant in the generator matrix and the rates $T^{\pm}_{M|\sigma}$ (this includes mutations) have a small effect on $P_M^{\rm st}$; if $m_g \gg \nu$, selective sweeps have small effect on $P_M^{\rm st}$ and the stationary distribution for $M$ is mainly determined by rates $T^{\pm}_{M|\sigma}$. When both effects are relevant, the prediction becomes less accurate (see central panels of Figures S8 and S9). In cases like this we cannot ignore the effect of selective sweeps on rates $T^{\pm}_{M|\sigma}$, neither we can ignore the effect of mutations on the stationary distribution.

As discussed in Section IV, for slow switching environments the distribution $P_M^{\rm st}$ is in general bimodal, with the lowest peak approaching $M = 1$ and increasing its probability as $\nu$ increases. This is also shown in Figure S8. The data in Figure S8 also show more clearly than in Figure 12 that the probability of the highest peak is not affected by $\nu$. For fast switching environments (see Figure S9), $P_M^{\rm st}$ is always unimodal with its mode approaching $M = 1$ as $\nu$ increases.

#### B. Fixed asexual environment

We now focus on a fixed environment with asexual reproduction, and explore how stationary distribution $P_M^{\rm st}$ behaves as parameters $m_g$ and $\nu$ are varied. This can be done by calculating the eigenvector with eigenvalue zero of a generator matrix given solely by the block $Q^{(A,A)}$ in Eq. (S94). The theoretical predictions and how they compare to numerical simulations in the full model are

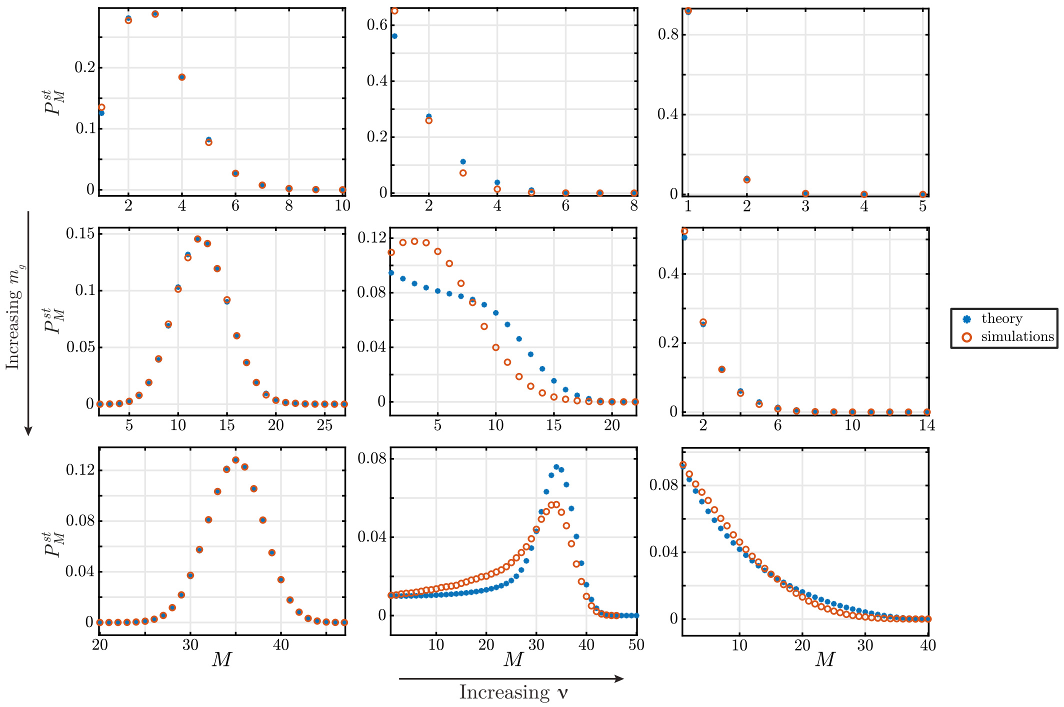


Figure S10. Theoretical prediction of $P_M^{\rm st}$ as function of $M$ for population size $N = 50$ and $\sigma = A$ (asexual environment) for different values of $m_g$ and $\nu$. From upper to lower rows: $m = 0.5, 5, 50$. From left to right columns: $\nu = 0, 0.01, 0.1$.

presented in Figure S10. As shown, the distribution is always unimodal for any value of $m_g$ and $\nu$, with the mode approaching $M = 1$ as $\nu$ increases. The fact there is only one peak is not surprising as the system has only one environment.

As stated in Section IV of the main paper, the mean time until fixation of a single mutant is approximately given by $T = 2\log(N)/s$, where $s$ the selective advantage of the mutant. Then, for a system that includes switching environments if we assume that $1/\lambda_{A\to S} \gg T$ , i.e., that the time spent in the asexual environment is long enough so that the beneficial mutation sweeps through the population, and that $p_S \approx 0$, i.e., that the fraction of time spent in the sexual environment is negligible (so we can ignore any effect of the sexual environment), we can estimate the corresponding stationary distribution $P_M^{\rm st}$ using only the block $Q^{(A,A)}$ as the generator matrix.


[1] George WA Constable and Hanna Kokko. The rate of facultative sex governs the number of expected mating types in isogamous species. *Nature Ecology & Evolution*, 2(7):1168, 2018.

[2] Silvia Heubach and Toufik Mansour. *Combinatorics of compositions and words*. CRC Press, Boca Raton, 2009.

[3] Melvyn B Nathanson. *Elementary methods in number theory*, volume 195. Springer Science & Business Media, New York, 2008.

[4] Steffen Eger. Restricted weighted integer compositions and extended binomial coefficients. *J. Integer Seq.*, 16(13.1):3, 2013.

[5] Lars Hörmander. *An introduction to complex analysis in several variables*. Elsevier, Amsterdam, 1973.

[6] PL Butzer and M Hauss. Riemann zeta function: Rapidly converging series and integral representations. *Applied mathematics letters*, 5(2):83–88, 1992.

[7] Ronald L Graham, Donald E Knuth, Oren Patashnik, and Stanley Liu. Concrete mathematics: a foundation for computer science. *Computers in Physics*, 3(5):106–107, 1989.

[8] Louis Comtet. *Advanced Combinatorics: The art of finite and infinite expansions*. Springer Science & Business Media, Dordrecht, 2012.
